# Sexual dimorphism in obesity is governed by RELMα regulation of adipose macrophages and eosinophils

**DOI:** 10.1101/2023.01.13.523880

**Authors:** Jiang Li, Rebecca E. Ruggiero-Ruff, Yuxin He, Xinru Qiu, Nancy M. Lainez, Pedro A. Villa, Adam Godzik, Djurdjica Coss, Meera G. Nair

## Abstract

Obesity incidence is increasing worldwide with the urgent need to identify new therapeutics. Sex differences in immune cell activation drive obesity-mediated pathologies where males are more susceptible to obesity co-morbidities and exacerbated inflammation. Here, we demonstrate that the macrophage-secreted protein RELMα critically protects females against high fat diet-induced obesity. Compared to male mice, RELMα levels were elevated in both control and high fat diet-fed females and correlated with adipose macrophages and eosinophils. RELMα-deficient females gained more weight and had pro-inflammatory macrophage accumulation and eosinophil loss, while both RELMα treatment and eosinophil transfer rescued this phenotype. Single cell RNA-sequencing of the adipose stromal vascular fraction was performed and identified sex and RELMα-dependent changes. Genes involved in oxygen sensing and iron homeostasis, including hemoglobin and lncRNA Gm47283, correlated with increased obesity, while eosinophil chemotaxis and response to amyloid-beta were protective. Monocyte-to-macrophage transition was also dysregulated in RELMα-deficient animals. Collectively, these studies implicate a RELMα-macrophage-eosinophil axis in sex-specific protection against obesity and uncover new therapeutic targets for obesity.

## Introduction

Obesity is an epidemic of significant public concern and contributes to the increased risk of several diseases, including type 2 diabetes, cardiovascular disease, nonalcoholic fatty liver disease, and COVID-19. Currently in the US, over 30% of men and women are classified as obese, with a body mass index (BMI) of ≥30kg/m^2^ (*1*). There are profound sex differences in adipose tissue deposition and obesity-associated diseases (*2*). Obese men are more at risk for metabolic syndrome, cardiovascular disease, and myocardial infarction than obese women (*3*). Male mice fed high fat diet (HFD) gain more weight and have an increased risk of insulin resistance than females (*4*). Despite these sex differences, most studies have historically focused on obesity mechanisms in males, since males gain weight more rapidly than females (*5*). Therefore, there remain many gaps in knowledge about the underlying mechanisms for obesity and whether these are sex-dependent, which can impact the development of therapeutics that are equally effective for both males and females. To address this gap, the focus of recent studies is identifying mechanisms that provide protection in females (*6, 7*). Males and females accumulate fat into different adipose tissue depots; males deposit more fat into visceral adipose depots, while females deposit fat preferentially into subcutaneous depots (*7, 8*). Since visceral adiposity is associated with the metabolic syndrome (*9*), differential fat accumulation may explain male propensity for obesity-mediated pathologies.

An underlying immune component for obesity pathogenesis is well recognized, with obesity being regarded as a chronic inflammatory process. Macrophages are critical immune effectors in obesity. Increases in adipose tissue size correlate with macrophage infiltration into the fat depots and proinflammatory cytokine production in both humans and mice (*10–12*). Obese adipose tissues produce increased levels of leptin (*6, 13*), and monocyte chemoattractant protein-1 (MCP-1, or CCL2 chemokine) that binds CCR2 (*14, 15*), which may serve as chemoattractant to recruit monocytes. In turn, they can be activated or differentiate into macrophages, initiating the secretion of cytokines and chemokines to exacerbate inflammation (*16, 17*). Given the role of macrophages in obesity, sex differences in macrophages are of particular interest and have been demonstrated before (*18, 19*). Visceral fat contains more infiltrating macrophages and higher expression of inflammatory cytokines than subcutaneous fat (*10, 20*), and male visceral adipose tissues accumulate more macrophages than females (*6, 21*). The presence of sex steroid hormones, specifically estrogen, was postulated to contribute to sex differences in obesity (*5, 8, 22*). An increase in adiposity following ovariectomy and removal of ovarian estrogen was observed in rodents (*23, 24*) and in monkeys (*25*), or after deletion of estrogen receptor alpha (*26*). However, we and others have demonstrated intrinsic sex-specific differences in macrophages, independent of sex-steroid hormones (*6, 27, 28*). Male macrophages are more migratory and inflammatory, while protection in females is associated with higher production of anti-inflammatory cytokines, such as IL-10 (*6, 7*). Macrophage function and activation in the adipose tissue are guided by their ontogeny, the cytokine environment, as well as myriad factors such as hypoxia, metabolites, lipids (*29*). CD11c^+^ M1 macrophages are activated through innate TLR2/4 receptors and produce proinflammatory mediators (e.g. TNFα, IL-6, CCL2) that drive metabolic changes. On the other hand, a T helper type 2 (Th2) cytokine environment within the adipose tissue promotes metabolic homeostasis and protective CD206^+^ M2 macrophages that suppress inflammation. Immune drivers of the Th2 cytokine environment for M2 macrophage activation include IL-4-producing eosinophils and innate lymphoid cell (ILC)-2 (*30, 31*). It is now recognized that macrophage activation is far more complex than the M1/M2 paradigm (*29, 32, 33*). However, the M1/M2 macrophage paradigm is a useful framework to begin to address pathways that can be targeted for obesity pathogenesis, and whether these are influenced by sex.

The focus of this study was to identify sex-specific immune effectors that regulate obesity pathogenesis, focusing on the M2 macrophage signature gene Resistin-like molecule α (RELMα). RELMα is a small, secreted cysteine-rich protein that is expressed by macrophages primarily in response to Th2 cytokines, but can also be induced by hypoxia (*34, 35*). RELMα has pleotropic functions ranging from inflammatory or immunoregulatory to microbicidal roles (*36–38*). Within the myeloid population, RELMα is preferentially expressed in monocyte-derived macrophages, and is important for monocyte differentiation, infiltration into other tissues and survival (*39, 40*). In the adipose tissue, RELMα is a defining marker for perivascular macrophages and is co-expressed with CD206 and Lyve1 (*29, 33*). A beneficial function for RELMα in metabolic disorders has been proposed; CD301b^+^ phagocytes promoted glucose metabolism and net energy balance through secretion of RELMα, and RELMα-overexpression promoted cholesterol homeostasis in hyperlipidemic low density lipoprotein receptor-deficient mice (*41, 42*).

In this study, we report a previously unrecognized sex-specific function of RELMα in protection from diet-induced obesity through promotion of eosinophils and regulation of adipose macrophage differentiation. Female mice expressed significantly higher serum and adipose RELMα than males, which correlated with increased adipose eosinophils, and protection from diet-induced obesity and inflammation. This protection was abrogated in RELMα-deficient (KO) females, which suffered from increased weight gain and adipose tissue inflammation, including greater proinflammatory macrophage infiltration. Eosinophil adoptive transfer and RELMα treatment was sufficient to rescue RELMα KO females from diet-induced obesity and inflammation. Single cell RNA sequencing (scRNA-seq) of the adipose stromal vascular fraction revealed cellular and molecular targets of female and RELMα-mediated protection, including PDGFR^+^ fibroblasts and ILC2, the kinase *Pim3*, and the long non-coding RNA *Gm47283*. Myeloid subsets were significantly impacted by sex and by RELMα deficiency; monocyte to perivascular macrophage differentiation was dysregulated in RELMα-deficient mice. Macrophages from female mice were enriched for eosinophil chemotactic factors, such as CCL24. Strikingly, oxygen-binding hemoglobin genes were highly upregulated in RELMα-deficient macrophages from female mice, which had enriched pathways associated with hypoxia and iron homeostasis. These data suggest a new mechanism for obesity pathogenesis, through macrophage-mediated oxygen level dysregulation and iron depletion. Taken together, we demonstrate a RELMα-eosinophil-M2 macrophage axis in female-specific protection and identify through scRNA-seq new targets for obesity including long non-coding RNAs and RELMα-regulated macrophage-specific hemoglobin genes.

## Results

### RELMα protects female mice from high fat diet-induced obesity and inflammation

Studies investigating sex differences show that female mice are protected, or have delayed, diet-induced obesity, unless aged or challenged by ovariectomy (*7, 43*). In support of the critical role of macrophage polarization in sex-specific differences, previous studies demonstrated that protection in females is associated with Th2 cytokine-induced M2 polarization, for example CD206 expression, while males exhibit increased CD11c-positive ‘proinflammatory’ M1 macrophages in the adipose tissue (*21*). The secreted protein RELMα is a signature protein expressed by M2 macrophages, with regulatory functions in downregulating inflammation and promoting tissue healing. A role for RELMα in promoting metabolic homeostasis has also been reported (*41, 42*). Based on these studies, we hypothesized that female-specific protection from high fat diet (HFD) may be influenced by RELMα. To examine systemic and local factors that may provide protection to females, serum and visceral adipose tissue homogenate were obtained from male or female mice on a control-fed (Ctr) or high fat diet-fed (HFD) for over 12 weeks. Under both Ctr and HFD conditions, female mice had significantly higher RELMα in the serum and visceral adipose tissue than males (Fig 1A).

**Figure 1.**
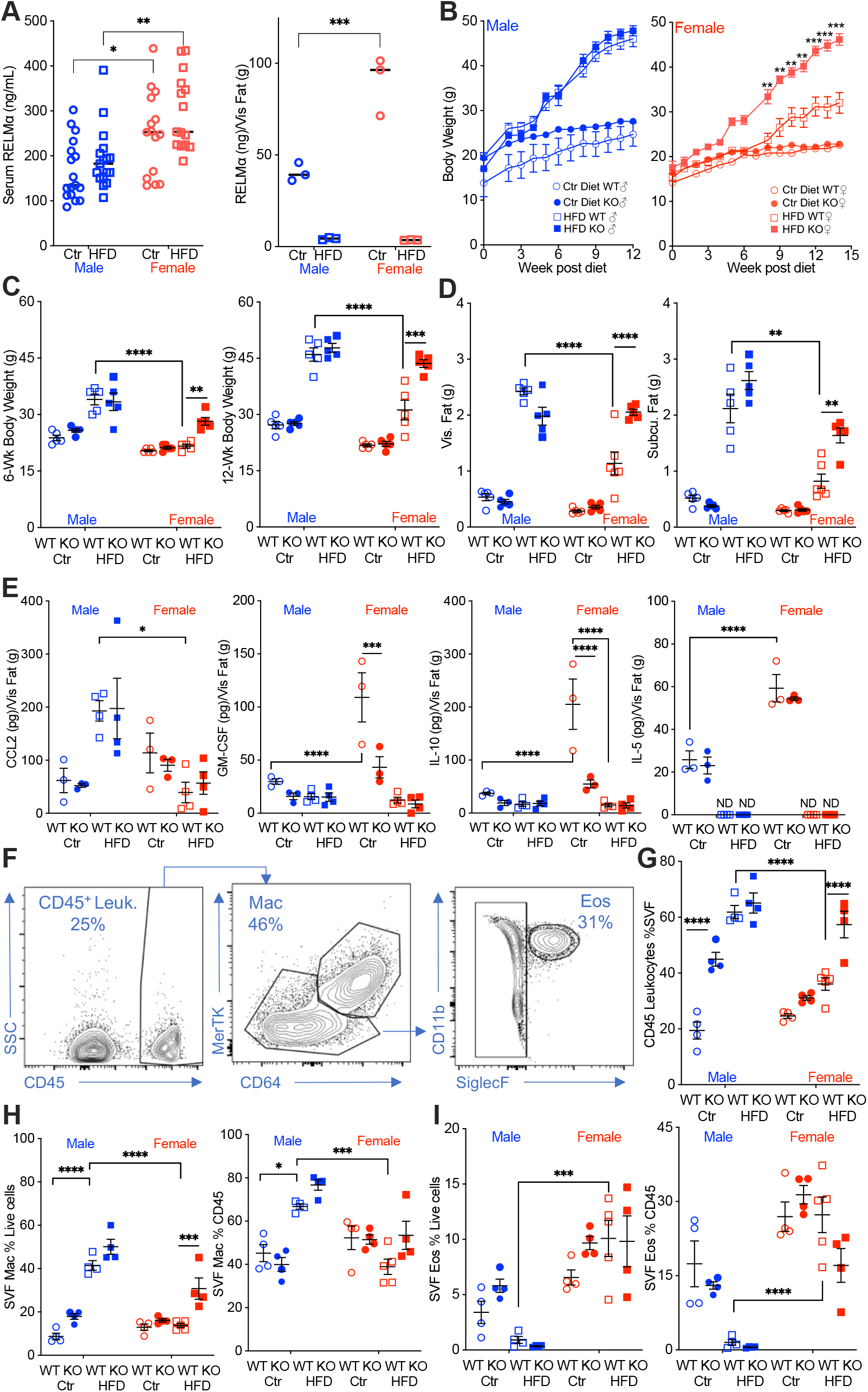
RELMα protects females from diet-induced obesity. **(A)** RELMα levels in serum and visceral adipose tissue from 18-week old male (♂) and female (♀) C57BL/6 mice after exposure to control (Ctr) or high fat diet (HFD) for 12 weeks. (**B**) WT or RELMα KO mice were weighed for 12-15 weeks of diet exposure. After 6-week diet and 12-week diet exposure (**C**), whole body, visceral and subcutaneous fat pad weights were recorded (**D**). (**E**) CCL2, GM-CSF, IL10, and IL5 levels in protein extracts from visceral fat pad after 12-week diet exposure. (**F**) Gating strategy for flow cytometric analysis of the visceral adipose stromal vascular fraction (SVF). (**G-I**) Proportion in the SVF of CD45^+^ leukocytes (**G**), CD45^+^CD64^+^Mertk^+^ macrophages (**H**), and CD45^+^SiglecF^+^CD11b^+^ eosinophils (**I**). Males (blue), females (red), WT (open symbols), RELMα KO (filled symbols), control diet (Ctr, circles), high fat diet (HFD, squares); data for (**B**) is presented as mean +/- S.E.M., data for (H) is representative of one animal, all other data are presented as individual points for each animal, where lines represent group means +/- S.E.M. Statistical significance between HFD WT females and HFD RELMα KO females was determined by unpaired t-tests for (B); and two or three-way ANOVA with Sidak’s multiple comparisons test for all other data. (ND, not detected; *, p < 0.05; **, p<0.01; ***, p<0.001; ****, p<0.0001 are indicated for functionally-relevant comparisons). Data are representative of 3 experiments with 4-6 mice per group.

To determine the role of RELMα in obesity, we placed RELMα knockout (KO) mice on Ctr and HFD, and compared their response to matched wild-type (WT) controls. RELMα deficiency did not affect Ctr or HFD weight gain in males, however, RELMα deficiency in females led to significantly increased weight gain on HFD compared to WT on HFD (Fig 1B). Whole-body weight, visceral and subcutaneous adipose weights were similarly increased in WT and KO males on HFD (Fig 1C-D). However, RELMα deficiency only affected HFD-fed females, with significantly increased body weight, and visceral and subcutaneous adipose tissue mass compared to HFD WT females. Chemokines that change with exposure to HFD were assessed in the adipose tissue of these mice (Fig 1E). The monocyte chemoattractant CCL2 was significantly elevated in HFD-males, while it remained low in females regardless of diet. On the other hand, females had higher levels of the anti-inflammatory IL-10 than males, as demonstrated before (*6, 7*), the regulatory T cell growth factor GM-CSF, and the Th2 cytokine IL-5. The higher level of IL-10 and GM-CSF in females were dependent on RELMα and were further decreased with exposure to HFD, while IL-5 was only decreased with HFD. Combined, the adipose protein profile indicates sex and RELMα-dependent effects of diet-induced obesity, which correlates with increased proinflammatory CCL2, and decreased anti-inflammatory and Th2 cytokines, IL-10, GM-CSF and IL-5, respectively.

Adipose tissue inflammation was next examined by flow cytometry of the stromal vascular fraction (SVF) from the visceral adipose tissue (Fig. 1F). RELMα deficiency and HFD resulted in significantly increased leukocyte frequency in the male SVF, demonstrating a role of RELMα in males (Fig 1G). WT females did not exhibit increased adipose tissue inflammation with HFD, as shown previously (*6*). Compared to HFD-fed WT females, RELMαKO females fed HFD had significantly increased SVF leukocytes, specifically macrophages (Fig 1H). In contrast, the proportion of eosinophils was decreased in HFD-fed males compared to females with the same diet (Fig 1I), suggesting a reduction in the protective type 2 immune response. This was consistent with the reduction in IL-5 in the adipose tissue following high fat diet (see Fig 1E). The contribution of diet, sex and genotype to body weight and adipose tissue inflammation was assessed by 3-way ANOVA (n=4-5 per group) (Supplemental Table 1). Diet, followed by sex then genotype, were all significant factors accounting for the variance in body weight, at both 6 and 12-weeks post diet. While diet was the greatest factor in adipose tissue inflammation, evaluated as SVF leukocyte frequency, RELMα deficiency was a greater factor accounting for variance than sex. For SVF macrophage frequency, diet then sex were the significant factors accounting for variance, while for eosinophil frequencies, sex differences were the main driving factor. Together, these data show that female-specific protection from diet-induced obesity is associated with elevated RELMα expression, and that RELMα deficiency selectively affects females, leading to increased weight gain, adipose tissue mass and adipose tissue inflammation.

### RELMα deficiency results in dysregulated macrophage activation and impaired eosinophil homeostasis in the adipose tissue

We employed t-distributed stochastic neighbor embedding (tSNE) analysis to evaluate immune cell heterogeneity and surface marker expression in the visceral adipose SVF (Fig. 2A). Within the groups, eosinophils demonstrated the greatest changes; Ctr-fed male and female mice had high eosinophil numbers, and this subset disappeared in male mice upon HFD (Fig. 2B, red outline). WT female mice retained their eosinophil subset even with HFD. In contrast, RELMα KO females had decreased eosinophils with HFD. Within the eosinophil subset, heterogeneity is observed in females, with Ctr mice exhibiting a different cell distribution compared to the HFD mice. Macrophage population also exhibited changes; Ctr-fed female mice, had a diminished macrophage subset compared to males, regardless of genotype (Fig. 2B, green outline). In both sexes, HFD led to an increase in this macrophage subset, and in their heterogeneity, especially in RELMα KO mice. Within the CD64^+^MerTK^+^ macrophage subset, expression of CD11c, a marker for proinflammatory M1 macrophages, was evaluated. There was an increase in both number and surface expression on a per cell basis of CD11c on macrophages in response to HFD in both males and females, however males had higher levels of CD11c than females under both diet conditions (Fig. 2C). RELMα deficiency exacerbated the increase in CD11c, particularly in the HFD-fed females (Fig. 2D). This may indicate that increased CD11c arises due to the abrogation of RELMα levels in the visceral fat after HFD (see Fig. 1A). On the other hand, anti-inflammatory ‘M2’ macrophage marker, CD206, decreased with HFD, but it was not dependent on the presence of RELMα (Fig. 2E). CD301b, another M2 marker that is upregulated by IL-4, also decreased with HFD in both males and females. However, specifically in HFD females, CD301 was further decreased with a loss of RELMα (Fig S1A). Although number of eosinophils decreased in HFD-fed males of both genotypes, eosinophils in all groups had high expression of Siglec-F, which was further increased with HFD in RELMα KO females WT females (Fig. 2F). We evaluated if eosinophil surface marker expression changed based on sex, diet, and genotype (Fig. S1B-D). CXCR4 and MHCII expression was reduced following HFD in both WT and KO females but not males, which may account for the subset heterogeneity (see Fig. 2B). Overall, these data identify that sex-specific and RELMα-dependent protection against diet-induced obesity is associated with changes in adipose macrophages and eosinophils.

**Figure 2.**
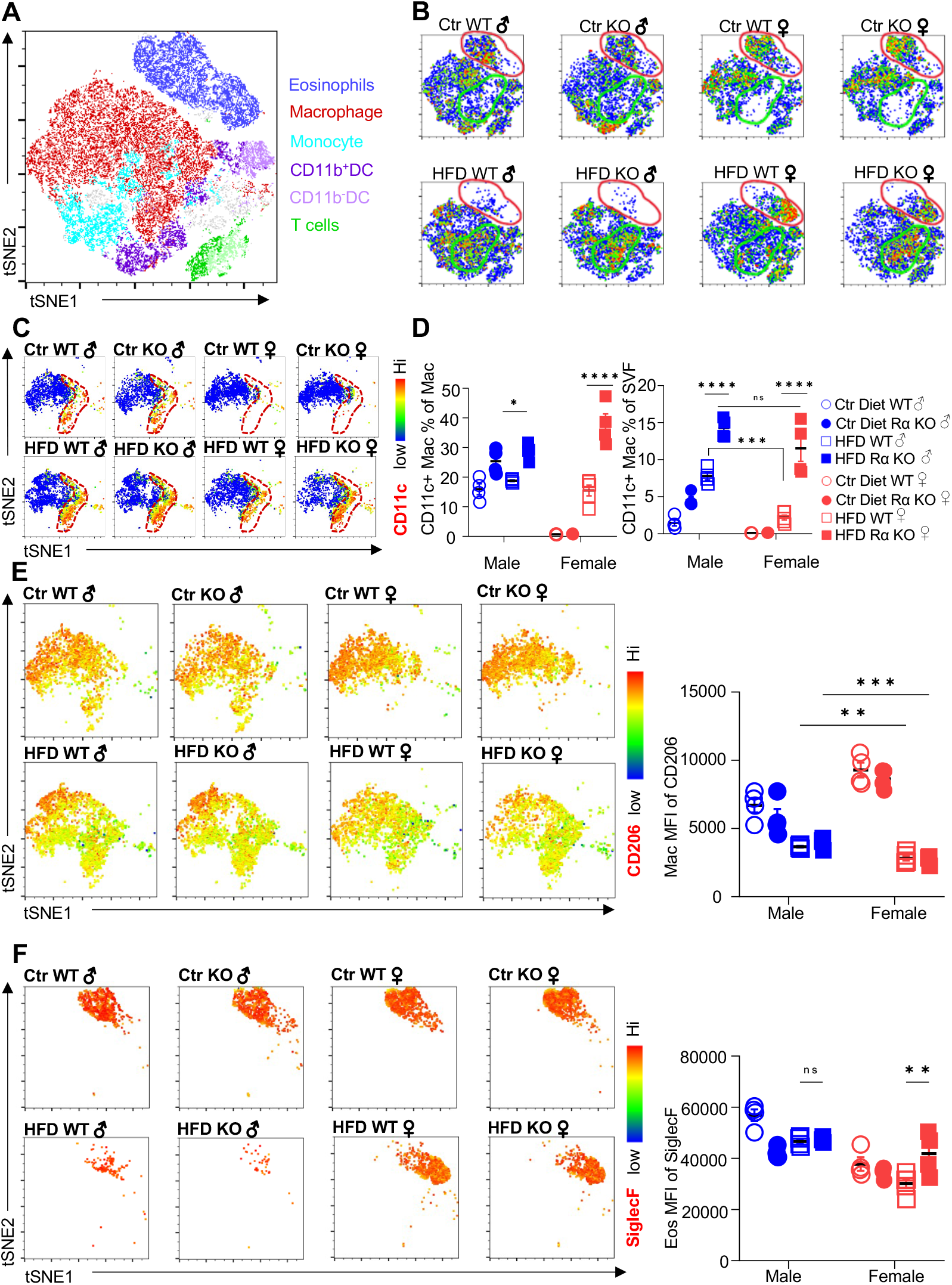
Adipose eosinophil and macrophage populations are influenced by sex, diet and RELMα. (**A**) tSNE analysis to identify SVF leukocyte populations. (**B**) tSNE analyses of SVF from the different groups (Male ♂, Female ♀, WT or RELMα KO) after 12 weeks of diet exposure (Ctr or HFD) revealed changes in eosinophil RELMα-dependent and diet-induced changes in eosinophil (red outline) and macrophage (green outline) subsets. (**C-D**) CD11c surface expression in CD45^+^MerTK^+^CD64^+^ macrophages was analyzed by tSNE, where dashed red outline shows CD11c^Hi^ cells, and quantified. (**E**) CD206 surface expression on SVF macrophages was examined by tSNE and quantified by mean fluorescent intensity (MFI). (**F**) Siglec-F surface expression on CD45^+^SiglecF^+^CD11b^+^ SVF eosinophils was examined by tSNE and quantified by mean fluorescent intensity (MFI). tSNE data are one representative animal per group. All other data are presented as individual points for each animal, where lines represent group means +/- S.E.M. Statistical significance was determined by three-way ANOVA Sidak’s with multiple comparisons test. (ns, no significant; *, p < 0.05; **, p<0.01; ***, p<0.001; ****, p<0.0001). Data are representative of 3 experiments with 4-6 mice per group.

### Protection against diet-induced obesity in females is mediated by RELMα and eosinophils

We evaluated whether associations existed between adipose immune cells and obesity by performing correlation analysis of body weight with adipose macrophage or eosinophil frequencies (Fig. 3A-B). Across mice from all groups, there was a significant, positive correlation between macrophage frequency and body weight in the visceral SVF. In contrast, SVF eosinophil frequencies were negatively correlated with body weight, implicating that adipose macrophage frequencies are linked to increased obesity while eosinophils are linked to lower body weight. RELMα expression has been reported by many immune cell subsets, including macrophages, eosinophils, B cells (*34, 44*), although expression in the adipose tissue is less clear. Given that RELMα protein was detectable in the visceral adipose tissue, especially of females (see Fig 1A), intracellular RELMα flow cytometry of the SVF cells was performed in Ctr or HFD-fed female mice. SVF macrophages expressed detectable RELMα, especially in the CD11c-negative subset, and this was significantly reduced with HFD (Fig 3C-D). RELMα^+^ SVF macrophage frequency was negatively correlated with body weight (Fig. 3E), supporting the protective role of ‘M2’ macrophages in obesity. Focused comparisons between HFD vs Ctr-fed female mouse groups showed that significant, negative correlation between RELMα^+^ macrophages and body weight only occurred with HFD-fed and not Ctr-fed mice, suggesting that the protective effect of RELMα may be impaired with body weight gain (Fig. 3F-G). Immunofluorescent staining of visceral adipose tissue sections was consistent with the flow cytometry analysis (Fig. 3H-I); HFD-fed male mice had increased F4/80^+^ macrophage crown-like structures (green), while HFD-fed females had fewer F4/80^+^ macrophages but had more detectable SiglecF^+^ eosinophils (magenta). In contrast, eosinophils and RELMα were absent from HFD-fed RELMα KO females, which had increased F4/80^+^ cells compared to HFD-fed WT females. These data implicate RELMα-driven eosinophils as the underlying mechanism of female-specific protection from HFD-induced obesity and adipose tissue inflammation. This hypothesis was tested by adoptive eosinophil transfer or recombinant RELMα treatment.

**Figure 3.**
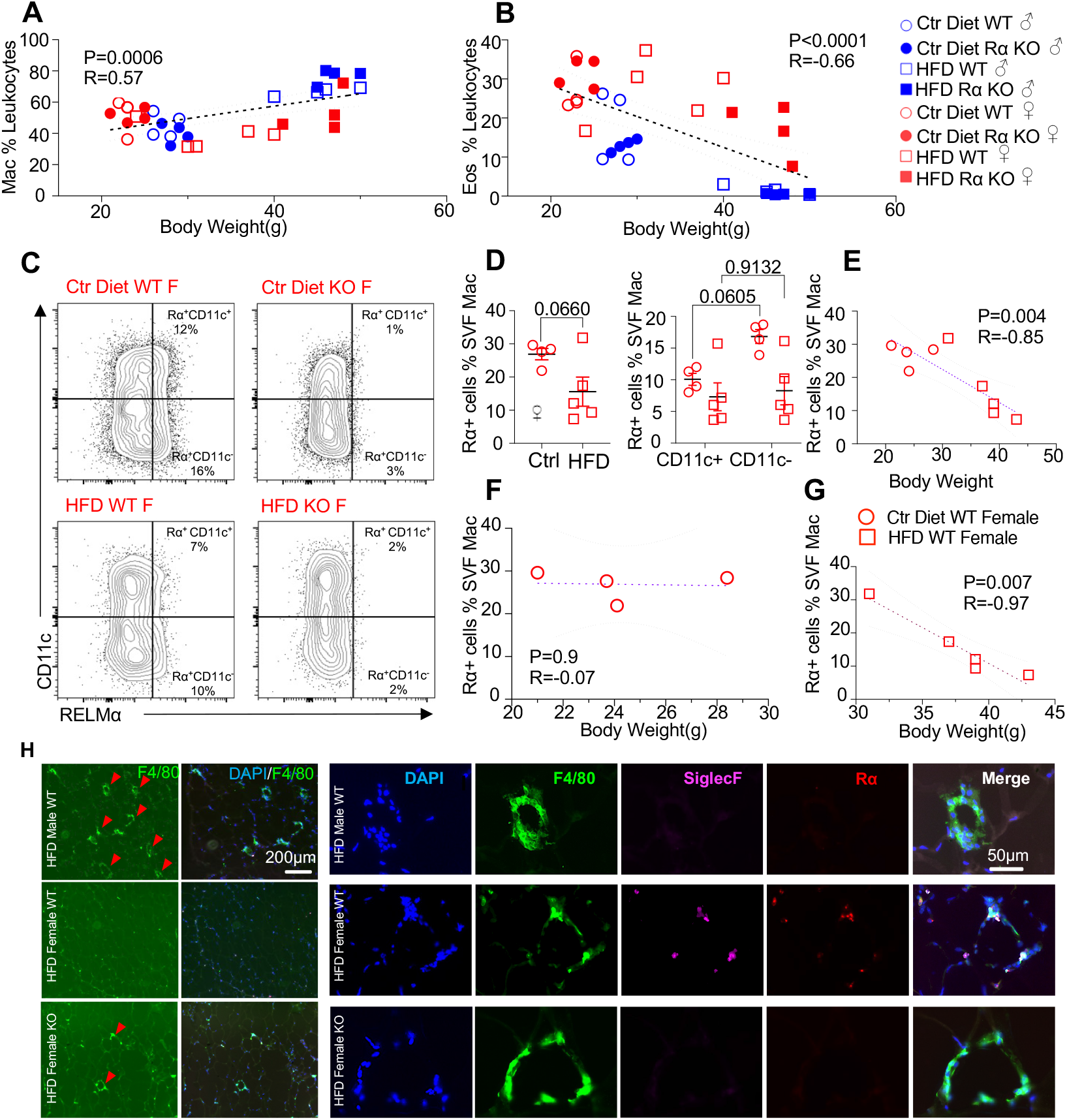
High fat diet-induced obesity is correlated with RELMα levels, eosinophils and macrophages. (**A-B**) Pearson correlation analysis of adipose SVF macrophage (**A**) or eosinophil (**B**) frequency against body weight of mice from all groups. (**C**) Representative flow plots of RELMα intracellular staining against CD11c surface staining of SVF Mac from WT and KO ♀ mice. (**D**) Frequency of RELMα^+^ SVF Mac in Ctr and HFD WT ♀ (left) or CD11c^+^ and CD11c^-^ Mac (right). (**E-G**) Pearson correlation analysis of RELMα+ cells against body weight of Ctr or HFD WT ♀ mice. (**H**) Immunofluorescent staining for F4/80 (green), SiglecF (magenta), RELMα (red) and DAPI (blue) was performed on visceral fat tissue sections (Bar, 200μM; red arrows indicate F4/80^+^cells). Flow plots (C) and IF images (H) are one representative animal per group. All other data are presented as individual points for each animal, where lines represent group means +/- S.E.M. Statistical significance was determined by unpaired t test (D), or Pearson correlation analysis for other data and p values are provided. Data are representative of 2 experiments with 4-6 mice per group.

Following previously published methodologies for eosinophil adoptive transfer to protect against obesity (*30, 31*), SiglecF^+^ eosinophils were column-purified from WT female mice that were chronically infected with helminth *Heligmosomoides polygyrus*, to increase eosinophil frequency (Fig. S1E-F). PBS or eosinophils (Eos) were intraperitoneally transferred into HFD-fed WT or RELMα KO female mice every 14 days, and weight gain monitored for 7 weeks, followed by analysis of the peritoneal and visceral adipose tissue. As an alternative approach, RELMα KO female mice were treated with recombinant RELMα with the same timeline. As expected, PBS-treated RELMα KO females gained significantly more weight than PBS-treated WT mice, however, this was rescued by both eosinophil adoptive transfer and RELMα treatment, with the KO+Eos and KO+RELMα having equivalent body weight to WT+PBS and WT+Eos (Fig. 4A-C). Flow cytometry analysis of the peritoneal cavity and visceral fat confirmed reduced eosinophils in RELMα KO compared to WT mice, which was rescued by eosinophil transfer or recombinant RELMα treatment (Fig. 4D-E). Evaluation of CD11c+ M1 macrophages in the visceral fat confirmed that RELMα KO mice had higher M1 macrophages compared to WT mice, but these were significantly decreased by either eosinophil transfer or RELMα treatment. These data identify a RELMα-eosinophil-macrophage axis underlying female-specific protection from diet-induced obesity and inflammation; macrophage production of RELMα is necessary to promote adipose eosinophil homeostasis and inhibit M1 macrophage activation, which is protective against HFD in females but not males.

**Figure 4.**
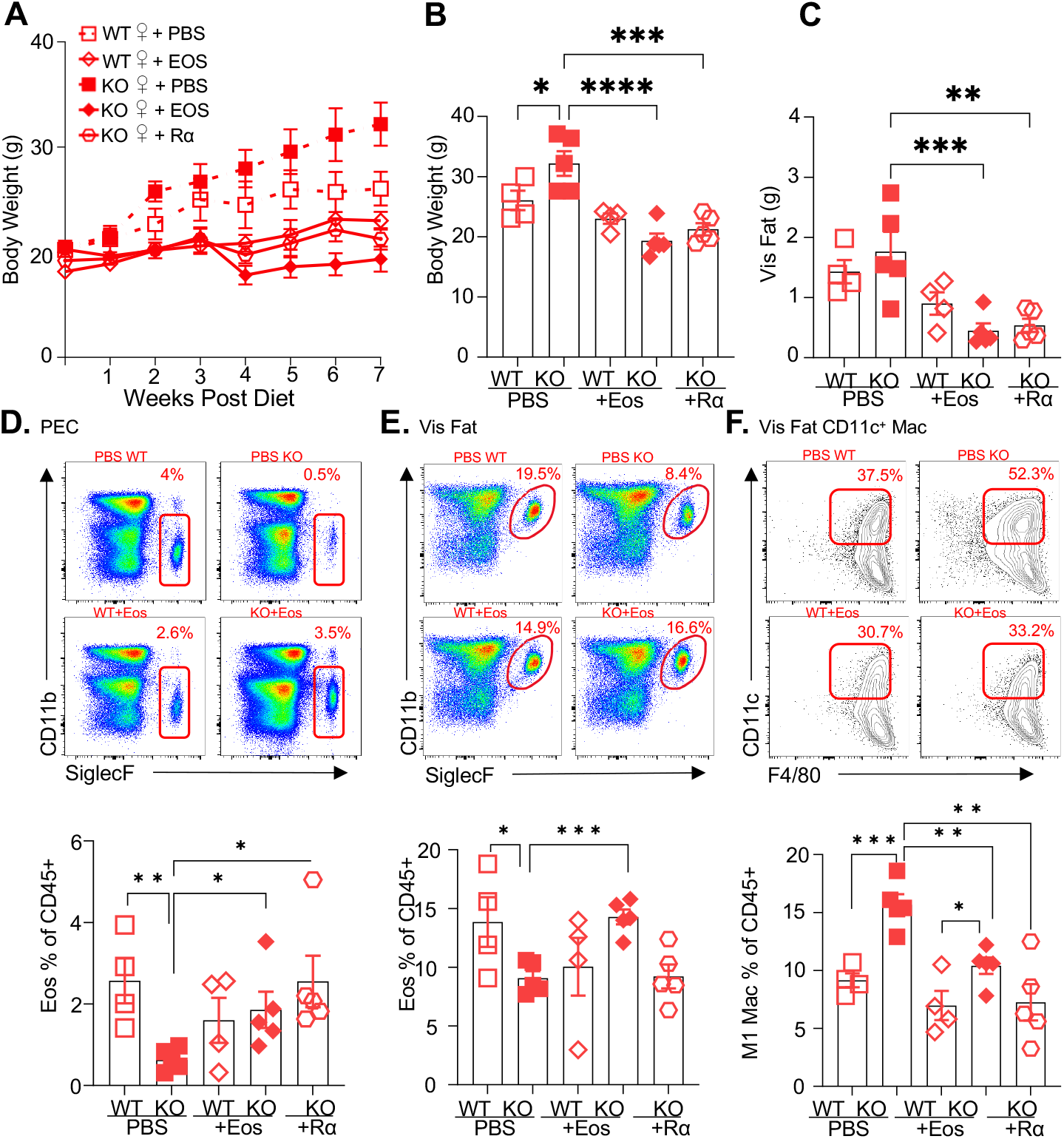
RELMα and eosinophils protect against diet-induced obesity. WT or RELMα KO female (♀) mice were exposed to HFD for 7 weeks, during which they were intraperitoneally injected every two weeks with PBS, RELMα (2μg) or SiglecF^+^ eosinophils (1×10^6^) recovered from helminth-infected WT ♀ mice. **(A)** Body weight was recorded every week. **(B-C)** Mice were sacrificed at 7 weeks post diet, and body **(B)** and visceral fat weight **(C)** were recorded. **(D-F)** Flow cytometric analysis and quantification of eosinophils from the peritoneal exudate cells (PEC) **(D)**, visceral fat SVF **(E)** and quantification of the CD11c^+^ Macs in the visceral fat SVF **(F)**. Data for **(A)** is presented as mean +/- S.E.M., flow plots for **(E-F)** are representative of one animal per group, all other data are presented as individual points for each animal, where lines represent group means +/- S.E.M. Statistical significance was determined by one-way ANOVA with Sidak’s multiple comparisons test. (ns, no significant; *, p < 0.05; **, p<0.01; ***, p<0.001; ****, p<0.0001). Data are representative of 2 experiments with 4-6 mice per group.

### Single cell RNA sequencing of the adipose stromal vascular fraction uncovers sex-specific and RELMα-specific heterogeneity

To identify cell-specific gene expression changes underlying RELMα-dependent and sex-dependent adipose effects, the 10x Genomics platform was used for single cell RNA-sequencing (scRNA-seq) of the visceral adipose SVF from 6-week HFD-fed WT *vs*. RELMα KO, and males *vs*. females. At 6-weeks HFD, WT females were protected from weight gain, compared to the other groups (see Fig. 1B), therefore this timepoint was chosen to define functionally relevant gene expression and pathway changes associated with weight gain. SVF single cell suspensions from each mouse per group were labeled with Cell Multiplexing Oligos (CMOs) to allow for pooling of biological replicates, prior to performing the single cell 3’ library generation and sequencing (Fig. 5A). Principal component analysis of all differentially expressed genes (DEG) confirmed clustering of biological replicates by group (Fig. 5B). A histogram of all DEG comparisons in all clusters between sex and genotype determined that WT male *vs*. WT female had the most DEG (Fig. 5C). Because WT females are protected from diet-induced changes, we sought to analyze gene expression changes in all clusters in WT females compared to WT males, KO females, and KO males (Fig 5D). A Venn diagram of the top 100 genes in WT females compared to WT males, KO females and KO Males revealed that WT females uniquely upregulated 75 genes compared to the other three groups (Fig. 5D). The enriched pathways in the top 75 genes that were upregulated in WT females compared to the three other groups were sex-specific (e.g. ovulation, sex differentiation, gonad, reproduction) (Fig. S2A-B). Non-sex specific pathways that were enriched in protected WT females included vasculogenesis and response to lipids, providing molecular hints to genes that are associated with protection from obesity and inflammation, e.g. TGFβ (Tgfb1, Tgfbr2, Tgfbr3), Th2 cytokine signaling (Il4, Il4ra, Il13) and matrix remodeling (connexins, metallopeptidases). A heatmap of the top 30 genes showed that WT females have increased expression of *Pim3*, a proto-oncogene, and anti-apoptotic gene *Bag3*, compared to the other three groups (Fig. 5E).

**Figure 5.**
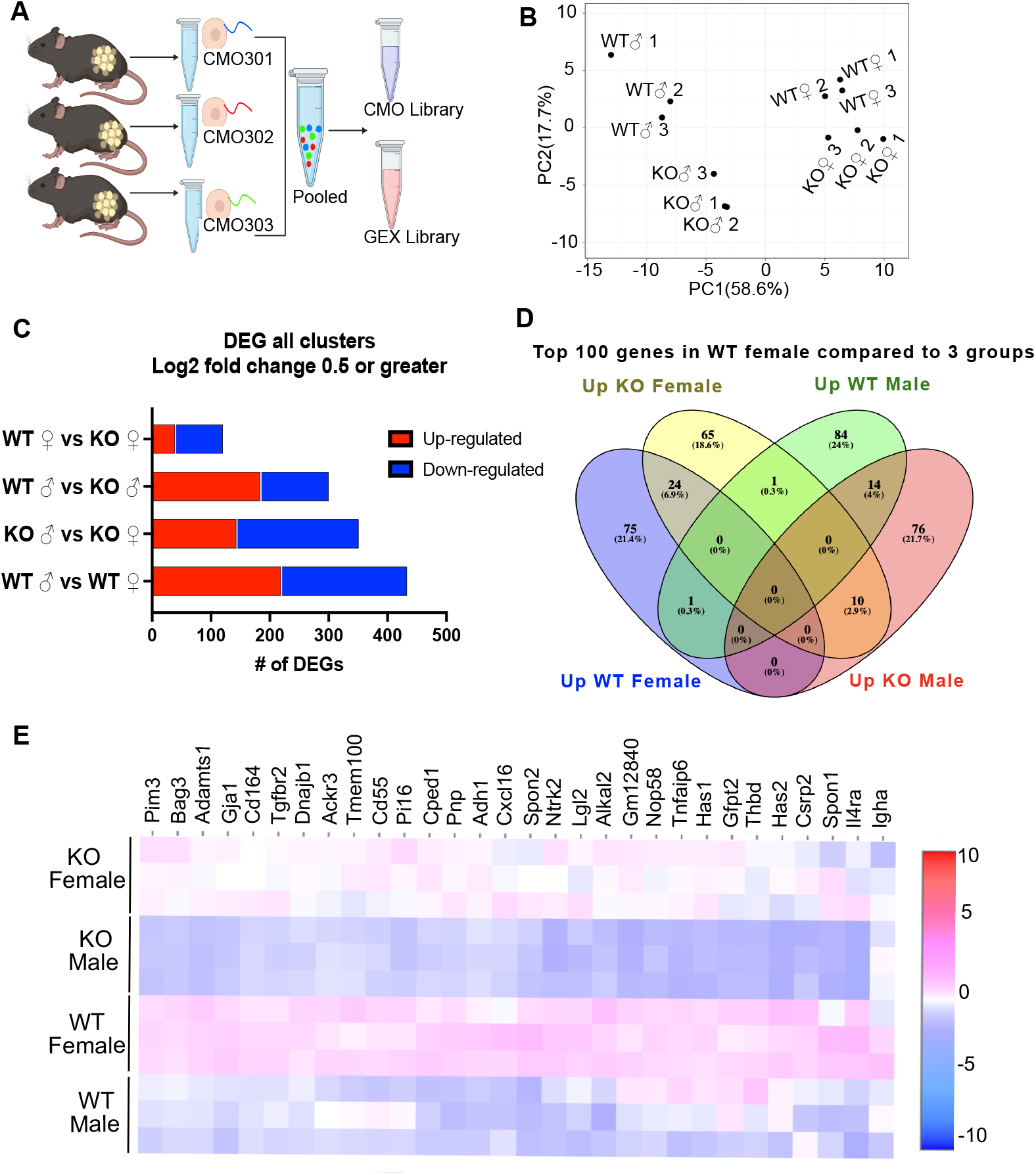
Single cell RNA-sequencing (scRNA-seq) of adipose stromal vascular fraction reveals genes associated with protection from diet-induced obesity in wild-type females. WT or RELMα KO male (♂) or female (♀) mice were exposed to HFD for 6 weeks, following which cells from the adipose stromal vascular fraction were recovered for single cell sequencing. **(A)** Schematic protocol of scRNA-seq cell multiplexing oligo (CMO) labeling and library preparation workflow. **(B)** Principal component analysis (PCA) assay of individual mice. **(C)** Histogram of differentially expressed gene (DEG) analysis with Log2 fold change of 0.5 or greater from all clusters comparing sex and genotype. **(D)** Venn diagram of the Top 100 genes in all clusters comparing WT female to KO female, WT male and KO male. **(E)** Heatmap of top 30 DEGs in WT female compared to all other groups in all clusters. Data are from 1 experiment with 3 mice per group.

The top differentially expressed genes and gene ontology (GO) pathways between males and females or WT and KO mice for all cells were examined (Fig. 6A-D). Comparison of WT females *vs*. WT males showed that most highly differentially expressed genes are as expected *Xist* (the X inactivation gene in females) and *Ddx3y* (unique to males, expressed on the Y chromosome). Other most highly differentially expressed genes upregulated in females include *Il4ra*, suggesting M2 macrophage responsiveness, and genes involved in extracellular matrix, such as *Spon2*, which promotes macrophage phagocytic activity (Fig. 6A). Males had higher levels of the sulfotransferase *Sult1e1*, and higher expression of inflammatory genes (e.g. *Lcn2*, lipocalin 2, and C7, complement 7) (Fig 6A). GO pathway analyses revealed that WT females upregulate genes in cellular responses to amyloid-beta. Compared to WT females, WT males upregulated genes in terpenoid and isoprenoid biosynthetic pathway, which are involved in cholesterol synthesis. Comparison between KO males and females revealed shared sex-specific DEG compared to WT mice (Fig 6B; upregulated *Xist, IL4ra*, and downregulated *Sult1e1* in KO females). However, unique sex-specific genes that were also identified were hemoglobin genes and oxygen binding pathways. These were upregulated in KO female mice compared to KO males. Macrophages can upregulate hemoglobin genes during inflammation and hypoxia (*45, 46*). On other hand, KO males upregulate genes involved in extracellular matrix (Collagen 4 genes) and vascularization (*Ccn2*).

**Figure 6.**
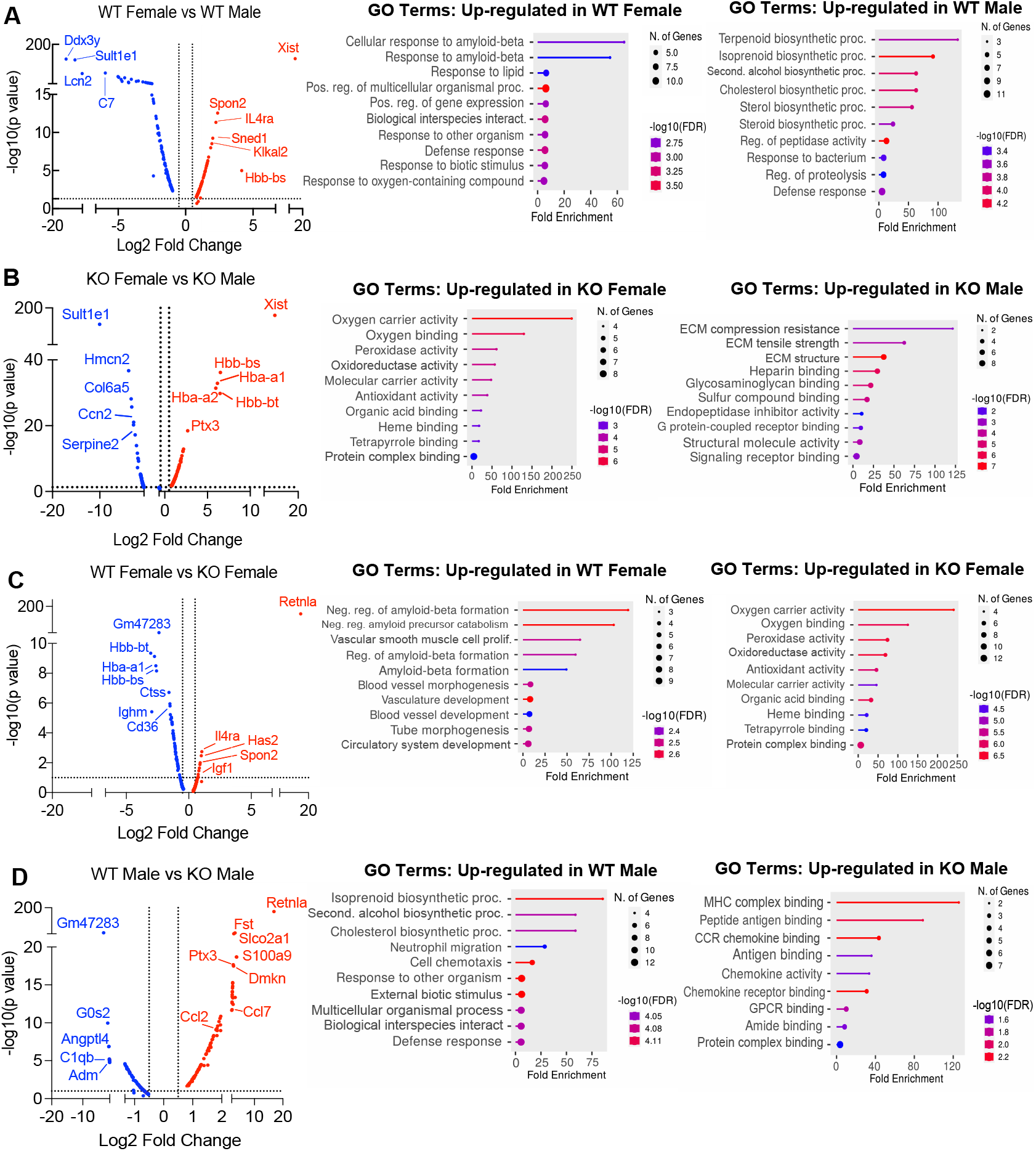
scRNA-seq identified sex-specific and RELMα-dependent gene expression and (GO) gene ontology pathway changes in diet-induced obesity. Volcano plot comparing the top 100 DEGs in all clusters between: WT females and. WT males **(A)**, KO females and KO males **(B)**, WT females and KO females **(C)**, WT males and KO males **(D)**. The most significant genes (-log10(p-value) >1, Log2 Fold change >0.5) are indicated. GO terms indicating enriched pathways for the top 30 upregulated genes are plotted as histograms. Data are from 1 experiment with 3 mice per group.

We then evaluated the RELMα-dependent genes that were associated with the loss of protection from diet-induced obesity in KO females (Fig. 6C). Genes related to the negative regulation of amyloid proteins were the most enriched pathways in WT females compared to KO females, following a similar trend to the WT female *vs*. WT male comparison. The downregulation of this pathway in WT males and KO females suggest that sex and RELMα contribute to protection through this shared pathway. On the other hand, RELMα KO females upregulated hemoglobin genes and oxygen-binding genes. Of note, these RELMα-driven differences were unique to females since they were not identified in the comparison between WT *vs*. KO males. Instead, RELMα-regulated genes in males mapped to innate inflammatory response pathways (e.g. increased genes related to MHC Class 2, chemokine/chemokine receptor signaling in the KO males), while genes in cholesterol synthesis pathway were upregulated in WT males compared to KO males (Fig. 6D). Of interest, long non-coding RNA *Gm47283* is the most upregulated gene in both male and female RELMα KO mice compared to their WT counterparts (Fig. 6C-D; Log2 fold change of 3.3 in KO males vs Log2 fold change of 2.4 in KO females). Together, these data suggest that female-specific genes regulated by RELMα map to non-immune but hypoxic and iron stress-related pathways (hemoglobin, oxygen binding, and ferroptosis).

### Cell-specific gene expression changes in fibroblasts, ILC2 and myeloid subsets correlate with sex-specific and RELMα-dependent protection against diet-induced obesity

Cell-specific gene expression changes were evaluated. Based on expression of known marker genes, twelve clusters were identified, consisting of immune and non-immune cells, displayed as tSNE plots and histograms (Fig. 7A-B). Eosinophils were not detected in any of the clusters. This was also shown in other SVF scRNA-seq studies, which concluded that eosinophils do not have sufficiently different transcriptomes from other leukocytes, or that there was a bias in the software, or technical difficulty such as degranulation that leads to RNA degradation, which precluded eosinophil identification (*47*). We employed targeted approaches to identify eosinophil clusters according to eosinophil markers (e.g. *Siglecf, Prg2, Ccr3, Il5r*), and relaxed the scRNA-Seq cutoff analysis to include more cells and intronic content, but still could not find eosinophils. We conclude that eosinophils may be absent due to the enzyme digestion required for SVF isolation and processing for single cell sequencing, which could lead to specific eosinophil population loss due to RNases or cell viability issues.

**Figure 7.**
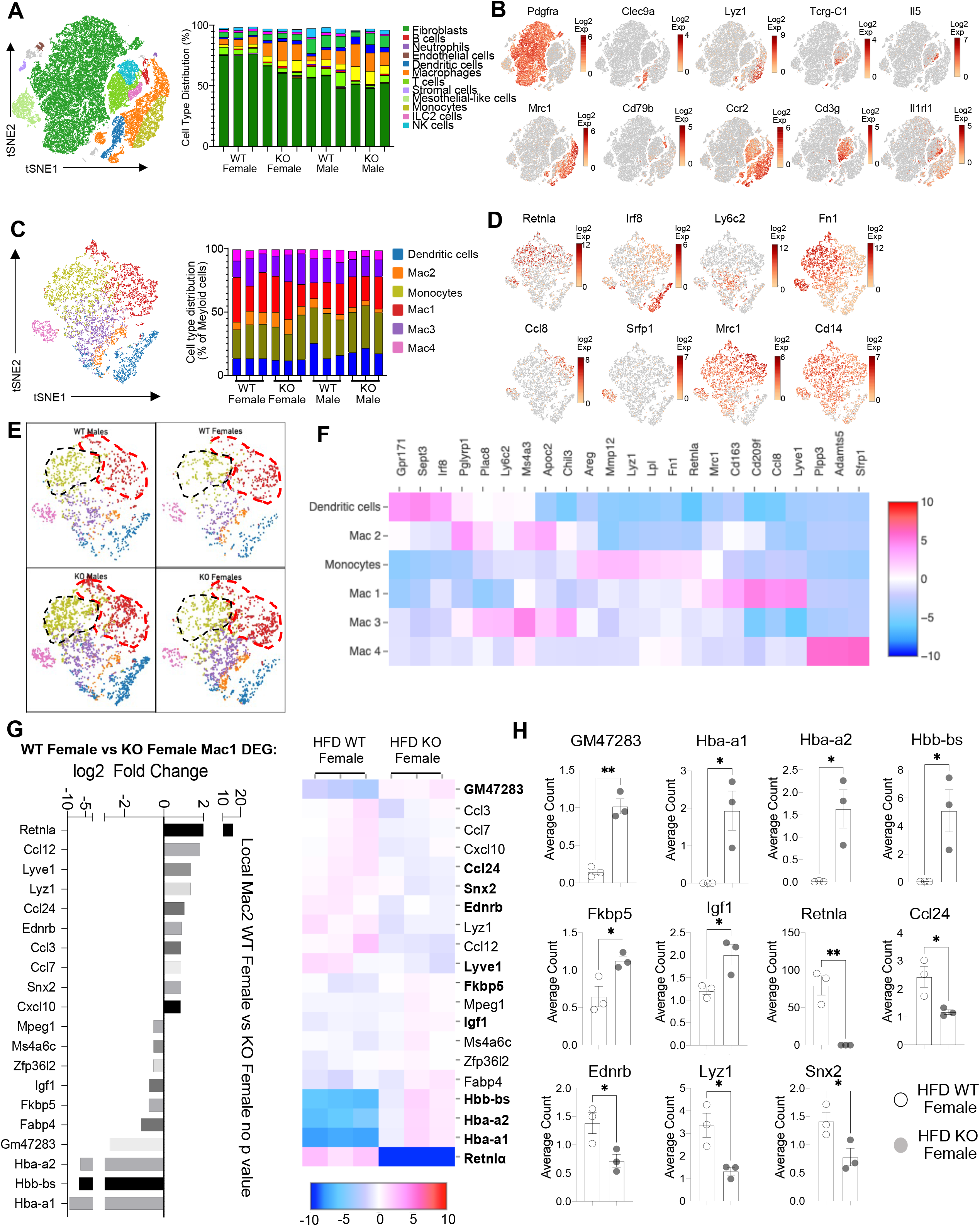
Sex-specific and RELMα-dependent gene expression changes in the SVF myeloid subsets in response to high fat diet. **(A)** tSNE plot showing cell populations from stromal vascular fraction (SVF) from all four groups of mice fed HFD for 6 weeks, with a histogram plotting cell type distribution per animal per group in all clusters. **(B)** Log2 fold change of candidate marker genes for each cell population across all clusters. **(C)** tSNE plot of re-clustered myeloid cell populations with a histogram plotting cell type distribution per animal per group. **(D)** Log2 fold change of candidate marker genes across myeloid cell populations. **(E)** tSNE plot highlighting population changes in Monocyte and Mac1 clusters between WT male, WT female, KO male and KO female in myeloid cells. **(F)** Heatmap of the top DEG that define each Mac subset. **(G)** WT female vs. KO female top DEG in Mac1 cluster. **(H)** Histograms of the average UMI count change of select candidate genes between WT female and KO female in Mac1 cluster. Data in **(H)** is presented as individual points for each animal, where lines represent group means +/- S.E.M. Statistical significance was determined by unpaired t-test (*, p < 0.05; **, p<0.01). Data are from 1 experiment with 3 mice per group.

The main population in the SVF, accounting for 50-75% of cells, were non-immune cells identified as *Pdgrfa*+ fibroblasts (green). These were significantly increased in WT females compared to the other groups (p<0.01, S2C). Compared to WT males, SVF fibroblasts from WT females upregulated pathways involved in inhibition of cell proliferation and M2 macrophage responses (e.g. IL-4R, Insulin growth-like factor, IGF-R) (Fig. S2D-F). On the other hand, WT male fibroblasts upregulated genes involved in vasculogenesis and extracellular matrix deposition, such as collagen genes and *Ccn2*, which contributes to chondrocyte differentiation. Similar trends were observed between WT and KO females, indicating again that RELMα deficiency results in females that are more similar to males by adipose tissue gene expression. KO females had increased levels of fatty acid binding proteins, *Fabp4* and connective tissue development, *Ccn3* and *Mmp3* (Fig. S2G-I). In males, WT males SVF fibroblasts increased expression of prostaglandin transporter, *Slco2a1* and stromal chemokine *Cxcl12*, while KO males upregulated pathways involved in the inhibition of the antioxidative functions, resulting in the accumulation of reactive oxygen species and oxidative stress, such as *Txnip* (Fig. S2J-L). Innate lymphoid cell (ILC)-2 are drivers of Th2 cytokine responses and are protective in obesity (*31, 48, 49*). ILC2 were present at small frequencies in the SVF in all groups (Fig. S2M). Gene expression analysis revealed that ILC2 from WT female mice expressed significantly higher Th2 cytokines (*Il13, Il5*) and *Csf2*, encoding for GM-CSF (Fig. S2N-P), which fits the increased adipose protein levels of IL-5 and GM-CSF in females (see Fig. 1). Functional pathway analysis revealed that fatty acid metabolism genes were upregulated in WT females compared to males. When comparing WT and RELMα KO female mice, there was a reduction in *Csf2* in the KO females compared to WT females. These data indicate that ILC2 in females are functionally distinct from males, and may contribute to the protective Th2 cytokine environment and metabolic homeostasis in the adipose tissue.

Myeloid cells/macrophages were the main immune cell subset that changed in the SVF in response to sex and RELMα deficiency; macrophage proportions were lowest in WT females, but expanded in the other groups (Fig 7A, orange). Gene ontology analysis revealed that the IGF pathway, chemokine and cytokine activity pathways were upregulated in WT female myeloid cells, while innate immune activation (e.g. TLR-4, RAGE receptor) and extracellular matrix remodeling were higher in WT males (Fig. S3A). This data matches macrophage polarization signatures where protective M2 macrophages produce and are responsive to IGF, while M1 macrophages respond to danger signals (e.g. LPS, RAGE). RELMα-dependent changes were observed in both females and males (Fig. S3B). Upregulated pathways in WT myeloid cells all involved innate chemokines and migration. Counterintuitively, downregulated pathways in WT compared to KO myeloid cells in females were associated with adipose tissue browning (e.g. brown fat cell differentiation, cold-induced thermogenesis), which are generally associated with protection from obesity. These data implicate RELMα in promoting innate immune cell migration and inhibiting adaptive thermogenesis. Overall, these scRNA-seq data identify sex-specific and RELMα-dependent changes in the adipose tissue that are associated with obesity-induced inflammation. Drivers of obesity included increased macrophages and innate immune activation. On the other hand, protection from obesity involved increased fibroblasts, Th2 cytokine expression by ILC2, and chemokine expression by myeloid cells.

### Monocyte to Mac1 macrophage transition and functional pathways are dysregulated in RELMα deficient mice

Based on previous studies, macrophage sub-clusters were defined and enumerated according to the Jaitin *et al*. study (*33*), and subset-specific gene expression was examined (Fig. 7C-E). Comparison of the genes unique to each myeloid subset *vs*. the other subsets showed that monocytes were enriched for *Lyz1*, while Mac1 were lymphatic vessel-associated macrophages expressing *Lyve1* (Fig. 7F). When analyzing scRNA-seq data from WT mice, RELMα (*Retnla*) is expressed in the Mono and Mac1 clusters (Fig. 7D, F). We evaluated cell-intrinsic effects of RELMα on these subsets. Gene ontology analysis revealed strong enrichment for genes involved in leukocyte migration in both the Mono and Mac1 subsets from WT females, specifically eosinophil chemotaxis (Fig. S4A). Evaluation of the top DEG indicated similar gene expression by sex rather than genotype (Fig. S4B). Focused chemokine analyses identified sex and genotype-specific eosinophil-recruitment chemokines. In particular, the eosinophil-recruiting chemokine *Ccl24* was significantly reduced in Mono and Mac subsets from KO females compared to WT females. We examined the Mac1 subset in females, to identify RELMα-regulated genes within this population that typically expresses RELMα under normal conditions (Fig. 7G, H). Hemoglobins were the most highly expressed genes in KO female Mac1 cells compared WT females (5-10 log2fold change). The hypoxia-induced lncRNA *Gm47283* was also upregulated in KO female Mac1. Together, these genes might indicate a response to hypoxia or oxidative stress in KO female Mac1 cells.

A trajectory analysis was performed to assess the relationships between the myeloid clusters, and whether they changed based on sex or genotype (Fig. 8A). In WT females, monocytes were the point of origin, leading to the generation of Mac1 subsets. Mac2 and Mac3 were related but separate clusters. Dendritic cells (DC) and Mac4 were even more distinct from monocytes suggesting that they are resident and not monocyte derived. These trajectories were similar in WT males. However, in KO males and females, the clusters were no longer distinct, and the Mac1 cluster was able to become Mac2/3 clusters, suggesting that loss of RELMα leads to dysregulated differentiation of monocytes to Mac1 or Mac2/3 subsets. We evaluated Mono to Mac1 transition in WT females and observed enriched pathways in IL-4 responsiveness and chemotaxis (Fig. 8B). In contrast, Mono to Mac1 transition in RELMα KO females involved proton transport and ATP synthesis pathways, suggesting dysregulated differentiation leading to metabolically active, inflammatory Mac1 subsets. Together, these data implicate a critical function for RELMα in myeloid cell function and differentiation in the adipose tissue. First, we uncover a RELMα cell-intrinsic mechanism whereby RELMα-expressing Mono and Mac1 cells mediate leukocyte recruitment, and Mac1 preferentially recruits eosinophils. Second, RELMα is necessary to drive functional Mac1 differentiation; in the absence of RELMα, Mac1 cells become metabolically active and increase their oxygen binding capacity by upregulating hemoglobin genes. Given that Mac1 in the normal setting are defined as the protective, vascular-associated, and anti-inflammatory subset, the loss of function of this myeloid population may be the underlying mechanism for increased inflammation in RELMα KO mice.

**Figure 8.**
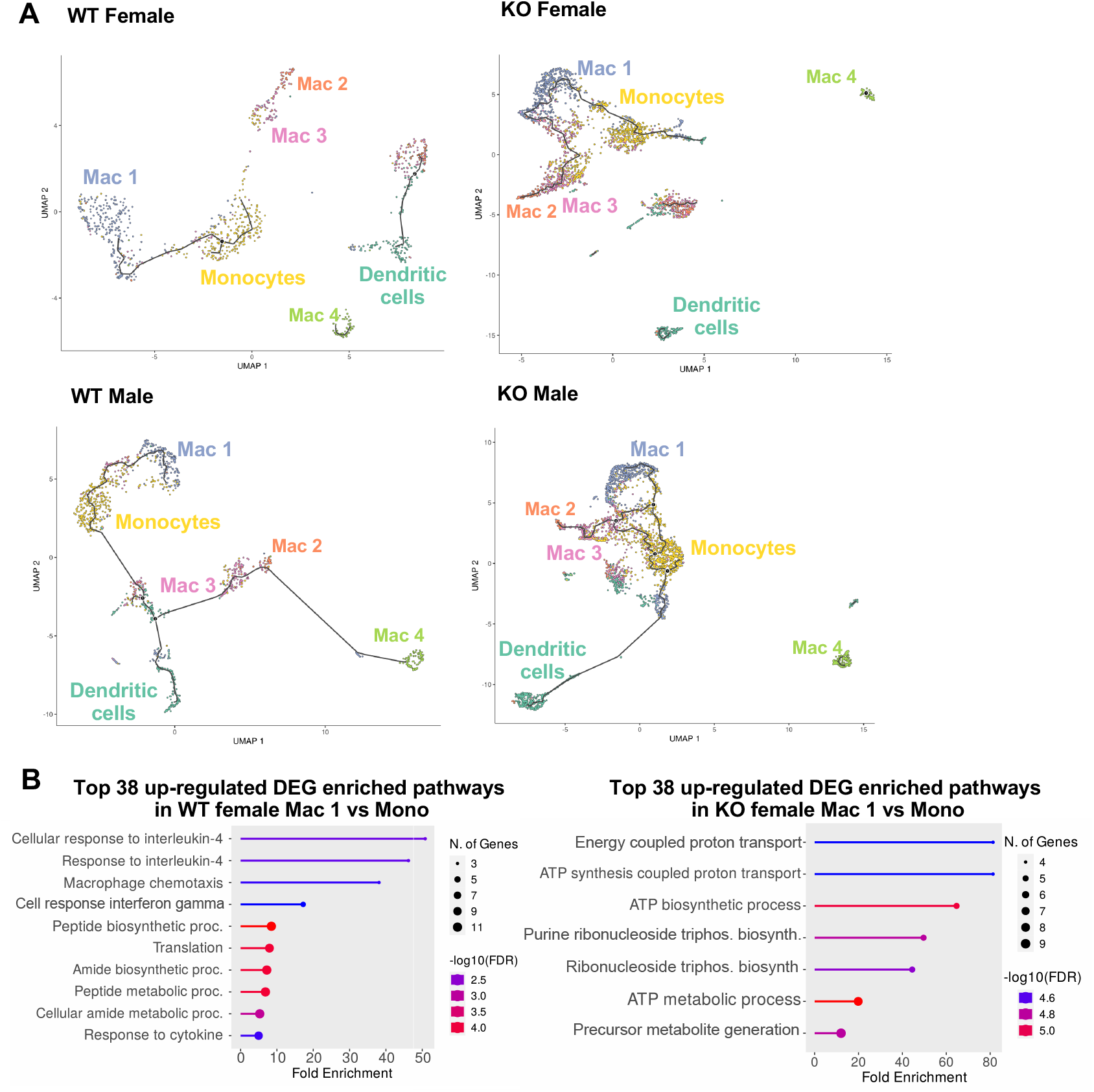
Trajectory analysis reveals dysfunctional myeloid differentiation in RELMα KO females. **(A)** UMAP plots of trajectory analysis with monocytes set as the root were made for the myeloid subsets within each group (WT female, WT male, KO female, and KO male fed HFD for 6 weeks). **(B)** Histogram of top 38 DEG with enriched GO terms that were upregulated in WT female Mac 1 vs. Monocyte population and KO female Mac 1 vs. Monocyte population. Data are from 1 experiment with 3 mice per group.

## Discussion

The goal of this study was to investigate sex differences in obesity pathogenesis and elucidate immune mechanisms underlying female-specific protection from adipose inflammation, cardiovascular disease, and the metabolic syndrome. Herein, we uncover an eosinophil-macrophage axis in females that is driven by RELMα and protects from diet-induced obesity and inflammation. A role for RELMα in whole body metabolism has been investigated (*41, 42, 50*), but none of these studies delineated sex-specific differences in chronic obese conditions. Here, by performing side-by-side comparisons between male and female RELMα KO or WT mice we identify sex-specific and RELMα-specific immune mechanisms of obesity pathogenesis. While C57BL/6J females are protected from obesity compared to males, we show that loss of RELMα abrogates this protection. RELMα deficiency also had significant effects in males, but to a lesser extent than females. For instance, RELMα KO males had increased proportions of leukocytes and CD11c^+^ macrophages in the stromal vascular fraction (SVF) to the same degree as exposure to high fat diet (HFD). Compared to WT females, RELMα KO females exhibited more diet-induced inflammatory changes than their male counterparts did. Under control and obese conditions, females had higher levels of RELMα than males, which likely explains why RELMα deficiency affected females more than males.

A significant strength of this study was the use of single cell sequencing to identify adipose tissue SVF heterogeneity and detect new targets and pathways to alleviate obesity. We first examined the top differentially expressed genes in protected WT females compared to the other groups and identify *Pim3* as protective. *Pim3* encodes a kinase that is a negative regulator of insulin secretion (*51*). *Pim3* is functionally responsive to forskolin, a cAMP activator that is also dietary supplement for weight loss and heart disease (*52, 53*). Our data indicate that forskolin’s effectiveness through Pim3 might be sex-dependent. As the most significantly upregulated gene in the protected WT females, our data also point to cAMP activation as a promising target to alleviate diet-induced obesity. Gene ontology (GO) pathway analyses revealed that WT females upregulate genes in cellular responses to amyloid-beta. Amyloid-beta synthesis is elevated in obesity in humans (*54*). In adipose tissue, it plays a role in lipolysis and secretion of adipokines (*55*). The increased cellular response to amyloid-beta specifically in females may explain female resistance to obesity-mediated changes. On the other hand, obese males had increased expression of *Sult1e1*, a sulfotransferase that leads to the inactivation of many hormones, including estrogen. *Sult1e1* is associated with increased BMI in humans (*56*). Males also had higher expression of inflammatory genes (e.g. *Lcn2*, lipocalin 2, and C7, complement 7). WT males also upregulated genes in terpenoid and isoprenoid biosynthetic pathway, such as *Aldh1a3, Fdps*, and *Hmgcs1* that regulate cholesterol synthesis, triacylglycerol absorption and fat deposition. Their association with insulin resistance and metabolic syndrome may explain the male propensity to develop these diseases (*57*). Examination of RELMα-dependent genes within the SVF led to the discovery of lncRNA Gm47283, which is the most significantly affected gene by RELMα deficiency. Very little is known about *Gm47283*, apart from one recent paper indicating it is a biomarker for myocardial infarction that is induced in hypoxia and involved in prostaglandin 2 synthesis and ferroptosis (*58*). Our findings indicate that RELMα potently downregulates this lncRNA, and future research is warranted to investigate whether Gm47283 is a downstream effector of RELMα.

Both control and HFD-fed females had a higher proportion of eosinophils in adipose tissues than males, and furthermore, males lost their eosinophil subset after exposure to high fat diet. Correlation analyses implicated that higher RELMα levels in females contributed to the higher proportion of eosinophils in female adipose tissues and female protection. This protection was lost in RELMα KO HFD-fed females, associated with the loss of eosinophils. Eosinophil transfer and RELMα treatment experiments confirmed this mechanistic link whereby RELMα recruits eosinophils with the overall outcome of reduced weight gain, decreased adipose tissue inflammation, and decreased CD11c^+^ proinflammatory macrophages. These findings support the potential for RELMα treatment in males to protect from obesity-mediated inflammation by driving eosinophils. A critical function for eosinophils in establishing a Th2 cytokine environment in the adipose tissue has been reported (*30, 59*). Specifically, through use of eosinophil-deficient mice or transgenic mice that have increased eosinophils, these studies demonstrate that eosinophils produce IL-4 to promote M2 macrophages, which in turn mediate adipose tissue beiging and other protective pathways against obesity. In the context of helminths, studies also identified eosinophils as the underlying mechanism whereby helminth infection protects from obesity. Our studies uncover further complexity to eosinophil function by demonstrating that females have significantly increased adipose eosinophils, and that eosinophilia is critically dependent on RELMα. Since females do not gain weight compared to males, investigation of female-specific pathways in murine models of obesity is an understudied area. However, there is an urgent need to determine what mechanisms are protective in females and whether these change with age or menopause. These would allow the identification of new therapeutic targets and will also distinguish whether treatments may differ in their effectiveness according to sex. Our findings open a new area of investigation into RELM proteins, which are produced in humans, and whether they regulate eosinophils to protect from obesity. Another study investigated whether IL-5-induced eosinophils could protect obese male mice from metabolic impairments but reported no protective effects (*60*). They concluded that physiological levels of eosinophils are not protective, in contrast to the previous studies that used transgenic mice to delete or artificially expand eosinophils. By additionally examining female mice and performing adoptive eosinophil transfer, our findings support a protective function for eosinophils even at physiologic levels, but also identify eosinophil heterogeneity. tSNE plot flow cytometric analysis of adipose SVF cells indicated that eosinophils were heterogeneous and different in females compared to males. These included changes in surface expression of CXCR4 and MHCII with HFD in females but not in males. In addition, single cell sequencing analyses of the adipose SVF indicated striking sex-specific or RELMα-specific changes in multiple eosinophil chemoattractants such as IL-5, produced by ILC2, and myeloid cell-derived eotaxin-2 (CCL24) and CXCL10. Eotaxin-2 was produced by myeloid cells in the SVF female WT mice, but was significantly decreased with the loss of RELMα. Our data implicates female-specific and RELMα-dependent immune mechanisms in the adipose environment, whereby ILC2 and myeloid cells recruit eosinophils that function to downregulate obesity-induced inflammation.

Myeloid cells are critical for adipose tissue homeostasis, and monocyte recruitment and differentiation to proinflammatory macrophages are associated with obesity. Strikingly, RELMα deletion led to induction of hemoglobin genes in SVF female KO mice compared to WT females and KO male mice. This may have significant health implications. The importance of hemoglobin in erythrocytes is well accepted, but the presence of hemoglobin in non-erythroid cells is less well known with limited studies. Hemoglobin gene induction was first detected in RAW264 and isolated peritoneal macrophages (*45*). Alternatively, hemoglobin genes can be induced by iron-recycling macrophages, derived from Ly6c^+^ monocytes during hemolysis, after erythrophagocytosis. Hemoglobin synthesis in cells other than erythroid lineage occurs in hypoxic conditions to increase oxygen binding and compensate for low oxygen (*61*). Therefore, it is possible that the lack of RELMα in females leads to hypoxia in adipose tissues. Alternatively, hemoglobin may be induced in KO females in response to macrophage activation and nitric oxide (NO) production, since hemoglobin can bind NO in addition to oxygen (*62*), which is produced by activated macrophages (*63*). Induction of hemoglobin genes may lead to dysregulation in iron handling and anemia, which have been associated with obesity. While obesity-increased incidence of anemia is not conclusive, iron deficiency is correlated with obesity (*64*). Macrophages normally recycle iron, but lack of RELMα in obese females may have disrupted this ability. Increase in hemoglobin gene expression may lead to iron sequestration, and would explain iron deficiency that is observed in obesity especially in women. Hemoglobin components include heme and iron, which can be cytotoxic. Overexpression of hemoglobin genes in the RELMα myeloid cells may not only act as a sink to deplete iron, oxygen and heme with consequences for the SVF environment, but could also constitute cytotoxic stress for the myeloid cells themselves, in a positive feedback cycle spurring further adipose dysfunction. To our knowledge, this is the first evidence of a hemoglobin pathway in myeloid cells during metabolic dysfunction and may point to new therapeutic targets and biomarkers for adipose tissue inflammation and obesity pathogenesis.

RELMα function in peritoneal macrophages was previously demonstrated to be sexually dimorphic, where peritoneal macrophage replenishment from the bone marrow is lower in females, and macrophage differentiation in females, but not males, is RELMα-dependent (*40, 65*). Our data further reveal that RELMα expression is sex-dependent and has critical functions in the adipose tissue through macrophage and eosinophil-driven mechanisms. We also demonstrate RELMα-specific effects on monocyte to macrophage transition in the adipose tissue that occur in both males and females. Whether these effects may be influenced by sex-specific differences in myeloid cell ontogeny from the bone marrow is unclear and an important avenue for future research. The importance of monocyte expression of RELMα for survival and differentiation has recently been reported (*66*). Trajectory analysis of the myeloid subsets revealed that WT animals of both sexes followed expected trajectories of monocyte differentiation to either Mac1 or to Mac2/3 clusters. Mac2/3 clusters express markers of proinflammatory macrophages, such as Ly6c, while Mac1 expresses markers of anti-inflammatory macrophages, such as *Mrc1* (CD206). The lack of RELMα in KO animals of both sexes led to dysregulated monocyte differentiation, where the ‘protective’ Mac1 cluster could become Mac2/3 cells. This trajectory change implies that lack of RELMα disrupts myeloid differentiation leading to a more proinflammatory profile. Genes enriched in monocyte to Mac1 transition in WT vs. RELMα KO female mice were examined to determine cell-intrinsic functions for RELMα. These analyses revealed that RELMα expression is critical for monocyte differentiation into IL-4 responsive macrophages, but in its absence, monocytes begin to increase expression of genes associated with high metabolic activity, which could result in oxidative stress.

In conclusion, these studies demonstrate a previously unrecognized role for RELMα in modulating metabolic and inflammatory responses during diet-induced obesity that is sex-dependent. Results from these studies highlight a critical RELMα-eosinophil-macrophage axis that functions in females to protect from diet-induced obesity and inflammation. Promoting these pathways could provide novel therapies for obesity pathology.

## Materials and Methods

### Animals

All experiments were performed with approval from the University of California (Riverside, CA) Animal Care and Use Committee (A-20210017; and A-20210034), in compliance with the US Department of Health and Human Services Guide for the Care and Use of Laboratory Animals. RELMα knockout mice were generated as previously described (*36*). RELMα and their WT controls were maintained under a 12-h light, 12-h dark cycle and received food and water *ad libitum*. After weaning and a week acclimatization on normal chow, animals were randomly distributed in groups and placed on either a high fat diet (HFD, D12492, 60% kcal from fat; 5.21 kcal/g (lard 0.32 g/g diet, soybean oil 0.03 g/g), 20% kcal from carbohydrate, 20% kcal from protein; Research Diet, New Brunswick, NJ) or control diet with matching sucrose levels to HFD (Ctr, D12450J, 10% kcal from fat 3.82 kcal/g (lard 0.02 g/g diet, soybean oil 0.025 g/g), 70% kcal from carbohydrate, 20% kcal from protein; Research Diet, New Brunswick, NJ) for 6-12 weeks, as indicated for each experiment.

### Eosinophil and RELMα treatment

For adoptive transfer, peritoneal exudate cavity cells were recovered from *H. polygyrus-infected* WT female mice (14-20 days post-infection). Eosinophils were column-purified with biotinylated anti-SiglecF (BioLegend), followed by anti-biotin MicroBeads then magnetic separation with MS columns according to manufacturer’s instructions (Miltenyi). 1×10^6^ eosinophils were transferred to recipient mice by i.p. injection. Eosinophil purity was confirmed by flow cytometry and by Diff-Quik stained-cytospins. For RELMα treatment, recipient RELMα KO female mice were i.p. injected 2 ug RELMα every 14 days. Ctr mice were injected with PBS.

### Cytokine Quantification

RELMα and IL5 were measured by sandwich ELISA. IFN-γ, CXCL1 (KC), TNF-α, CCL2 (MCP-1), IL-12p70, CCL5 (RANTES), IL-1β, CXCL10 (IP-10), GM-CSF, IL-10, IFN-β, IFN-α and IL-6 were detected by the Mouse Anti-Virus Response Panel (13-plex) (Cat# 740622 BioLegend, San Diego, CA) and analyzed on the NovoCyte Flow Cytometer (Agilent, Santa Clara, CA) and LEGENDplexTM software (Biolegend, San Diego, CA).

### Histological analyses and immunohistochemistry

At the conclusion of diet exposure, mice were anesthetized, perfused with 20 ml cold PBS, fat tissues were recovered and immersed in 4% PFA for 24 hours followed by 30% sucrose for another 24 hours. After, fat tissues were embedded with O.C.T (Sakura Finetek USA) and sectioned at 10 μm. For immunofluorescent staining, sections were incubated with APC-anti RELMα (DS8RELM, eBioscience, Santa Clara, CA), PE/Dazzle^™^ 594 anti-SiglecF (S17007L BioLegend, San Diego, CA) and Alexa Fluor^®^ 488 anti-mouse F4/80 (BM8 eBioscience, Santa Clara, CA) overnight at 4°C, then counterstained with DAPI (BioLegend, San Diego, CA). Slides were examined with the Keyence microscope (BZ-X800; lense:BZ-PF10P, Plan Fluorite 10X, WD 14.5 mm; BZ-PF40LP, Plan Fluorite 40X LD PH, WD 2.2 to 3.3mm).

### Flow cytometry

Tissues from each mouse were processed separately as part of a 3-4 mouse cohort per group, with each experiment repeated 2-3 times. In brief, mice were perfused with ice cold PBS, adipose tissue was collected from gonadal fat pads representing visceral fat depot, or from inguinal fat pads representing subcutaneous fat depot, rinsed in cold PBS, weighed, minced with razor blade and digested enzymatically with 3 mg/mL collagenase/dispase (Roche) at 37 °C for 1 hour. Suspension was passed through 70 μm cell strainer, cells pelleted, and red blood cells lysed using RBC lysis buffer (Biolegend, San Diego, CA). Stromal vascular fraction cells were collected and counted, and 2 million cells labeled for flow cytometry analyses. Cells were Fc-blocked with anti-mouse CD16/CD32 (1:100, Cat# 553141, BD Biosciences, San Jose, CA) followed by surface marker staining with antibodies to MerTK (2B10C42, Biolegend), CD25 (PC61, Biolegend), CD301 (LOM-14, Biolegend), CD36 (HM36, Biolegend), MHCII (M5/114.15.2, Biolegend), CD45 (QA17A26, Biolegend), CXCR4 (L276F12, Biolegend), CD11b (M1/70, Biolegend), F4/80 (BM8, Biolegend), CD4 (RM4-5, Biolegend), CD206 (C068C2, Biolegend), SiglecF (S17007L, Biolegend), CD11c (N418, eBioscience), CD64 (X54-5/7.1, Biolegend), RELMα (DS8RELM, eBioscience). Dead cells were labeled with Zombie Aqua^™^ Fixable Viability Kit (Cat# 423102 Biolegend, San Diego, CA). Gating strategy was followed: macrophage (CD45^+^CD11b^+^MerTK^+^CD64^+^), eosinophils (CD45^+^CD11b^+^SiglecF^+^ MerTK^-^CD64^-^), monocyte (CD45^+^CD11b^+^MHCII^+^CD11c^-^MerTK^-^CD64^-^SiglecF^-^), dendritic cells (CD45^+^ MHCII^+^CD11c^+^MerTK^-^CD64^-^SiglecF^-^) and T cells (CD45^+^ CD4^+^CD11b^-^MHCII^-^ CD11c^-^MerTK^-^CD64^-^SiglecF^-^). Cells were analyzed on the NovoCyte Flow Cytometer (Agilent, Santa Clara, CA) and FlowJo v10 software (Tree Star Inc.; Ashland, OR). tSNE analyses were performed using FlowJo v10 (Tree Star Inc.; Ashland, OR), following concatenation of samples (5000 cells per biological replicate) from all groups, to generate plots consistent between groups. This was followed by analysis of the expression of desired markers in separated groups. The parameters used to run the tSNE analyses were FITC-MerTK, PerCP-CD25, Alexa Fluor 700-MHCII, Brilliant Violet 605-CD11b, Brilliant Violet 650-F4/80, Brilliant Violet 711-CD4, PE/Dazzle 594-SiglecF, PE Cy5-CD11c and PE Cy7-CD64.

### Stromal vascular fraction (SVF) isolation from adipose tissue

Adipose tissue was dissected, rinsed in ice cold PBS, and minced with a razor blade. Fat was digested enzymatically with 3 mg/ml collagenase/dispase (Roche) at 37°C for 1 hour. Suspension was passed through a 70 μm strainer and centrifuged to pellet stromal vascular fraction. Cells were resuspended in PBS/0.04% BSA, counted, viability determined to be >90% before proceeding to flow cytometry analysis or scRNA-seq.

### Single cell RNA-seq (scRNA-seq)

ScRNA-seq was performed following the Chromium Next GEM Single Cell 3’ v3.1 Dual Index with Feature Barcoding for Cell Multiplexing protocol. Each group consisted of three biological replicates. SVF single cell suspension from each mouse was labeled with the 10X Genomics Cell Multiplexing Oligos (CMOs) following manufacturer’s protocol (10x Genomics, Demonstrated Protocol, CG000391). Cell suspensions from mice in the same group were pooled and processed for the Chromium Next GEM Single Cell 3’ v3.1 Dual Index (10X Genomics, Demonstrated Protocol, CG000388). 50,000 cells were loaded on the Chromium Chip for GEM generation and barcoded, in order to reach a targeted cell recovery of 30,000 cells per group. Following post-GEM cleanup, libraries were amplified by PCR, after which the library was split into two parts: one part for generating the gene expression library and the other for the multiplexing library. Libraries were indexed for multiplexing (Chromium Dual Index Kit TT Set A, PN-1000215), quantified by Qubit 3 fluorometer (Invitrogen) and quality assessed by bioanalyzer (Agilent). Equivalent molar concentrations of libraries were pooled and sequenced using Novaseq 6000 (Illumina) at the UC San Diego (UCSD) Institute for Genomic Medicine (IGM) Center.

### Data processing and analysis

Raw scRNA-seq data were demultiplexed and processed using the Cell Ranger v6.1.2 software (10x Genomics) with the multi pipeline and aligned to the Cell Ranger mm10-2020-A mouse reference genome, introns excluded. T-distributed stochastic neighbor embedding (tSNE) clustering and differential gene expression analysis were performed using Loupe Browser v6.2.0 (10x Genomics). Cells with threshold UMI count of >40,000 and <1000 and mitochondrial fraction of >10% were filtered from analysis. Gene ontology (GO) enrichment analysis of the genes from DEG analysis was performed using the ShinyGo 0.76.3 platform (South Dakota State University (*67*)). False discovery rate (FDR) cutoff was 0.05, with the pathway minimum set to 10. Trajectory analysis was performed using Monocle 3 pipeline (*68*) using single-cell batch correction methods and Seurat v3.9.9.9024 (*69*). Top markers for each cluster were compared to known markers of immune cell types to annotate the clusters.

## Data availability

The raw data have been deposited at the Gene Expression Omnibus, GEO accession number GSE219119 and is fully available at https://www.ncbi.nlm.nih.gov/geo/query/acc.cgi?acc=GSE219119

## Code availability

Experimental protocols and the data analysis pipeline used in our work follow the 10X Genomics and Seurat official websites. The analysis steps, functions, and parameters used are described in detail in the Methods section.

## Statistical analyses

Data are presented as mean ± SEM and statistical analysis was performed by Graphpad Prism 9. Statistical differences in biological replicates between control and RELMα knockout mice (p < 0.05) were determined using t-test, or 2-way or 3-way ANOVA with Sidak multiple comparisons test. *, p ≤ 0.05; **, p ≤ 0.01; ***, p ≤ 0.001 and **** p ≤ 0.0001. 10x scRNA-seq experiment was performed once (3 mice per group). All other *in vivo* experiments were repeated 2-4 times with n=3-5 per group (combined n=6-20). No outliers were excluded. Sample sizes were determined based on previous studies, and no statistical methods were used to predetermine sample size.

## Acknowledgements

We thank Brandon Le for bioinformatics advice; Dr. Karine Le Roch for access to the 10X single cell sequencing equipment; Constance Finney, Sang Woo and Sang Yong Kim for provision of the *Heligmosomoides polygyrus* larvae.

## Supplemental figures

**Figure S1.**
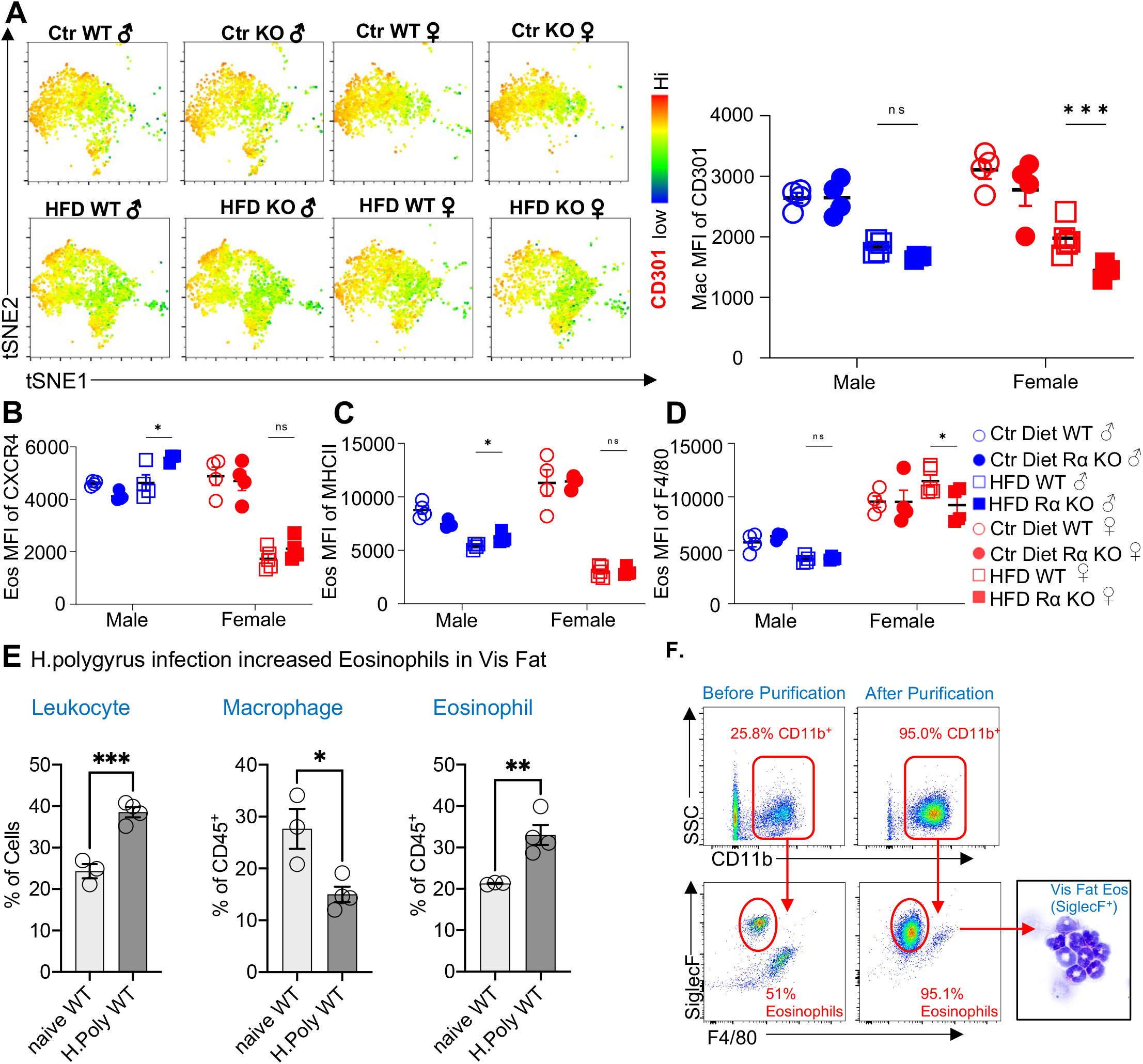
Flow cytometric analysis of adipose immune cells. **(A-D)** Flow cytometry analysis of the visceral adipose stromal vascular fraction of 12-week diet exposed mice for CD301 surface expression on CD45^+^CD64^+^Mertk^+^ macrophages by tSNE plot and mean fluorescent intensity (MFI) **(A)** and surface expression by CD45^+^SiglecF^+^CD11b^+^ eosinophils of CXCR4 **(B)**, MHCII **(C)** and F4/80 **(D)**. (**E-F)** Visceral fat leukocytes from naïve or *H. polyrus-infected* mice were quantified **(E)**, and eosinophils were column purified and analyzed for purity by flow and cytocentrifuge **(F)**. Statistical significance was determined by unpaired t-test (*, p < 0.05; **, p<0.01). tSNE plot analysis and flow plots are representative of one animal per group, all other data are presented as individual points for each animal, where lines represent group means +/- S.E.M. Statistical significance for (A-D) was determined by two-way ANOVA with Sidak’s multiple comparisons test. (ns, no significant; *, p < 0.05; **, p<0.01; ***, p<0.001) and by unpaired t-test for (E). Data are representative of two experiments with 3-6 mice per group.

**Figure S2.**
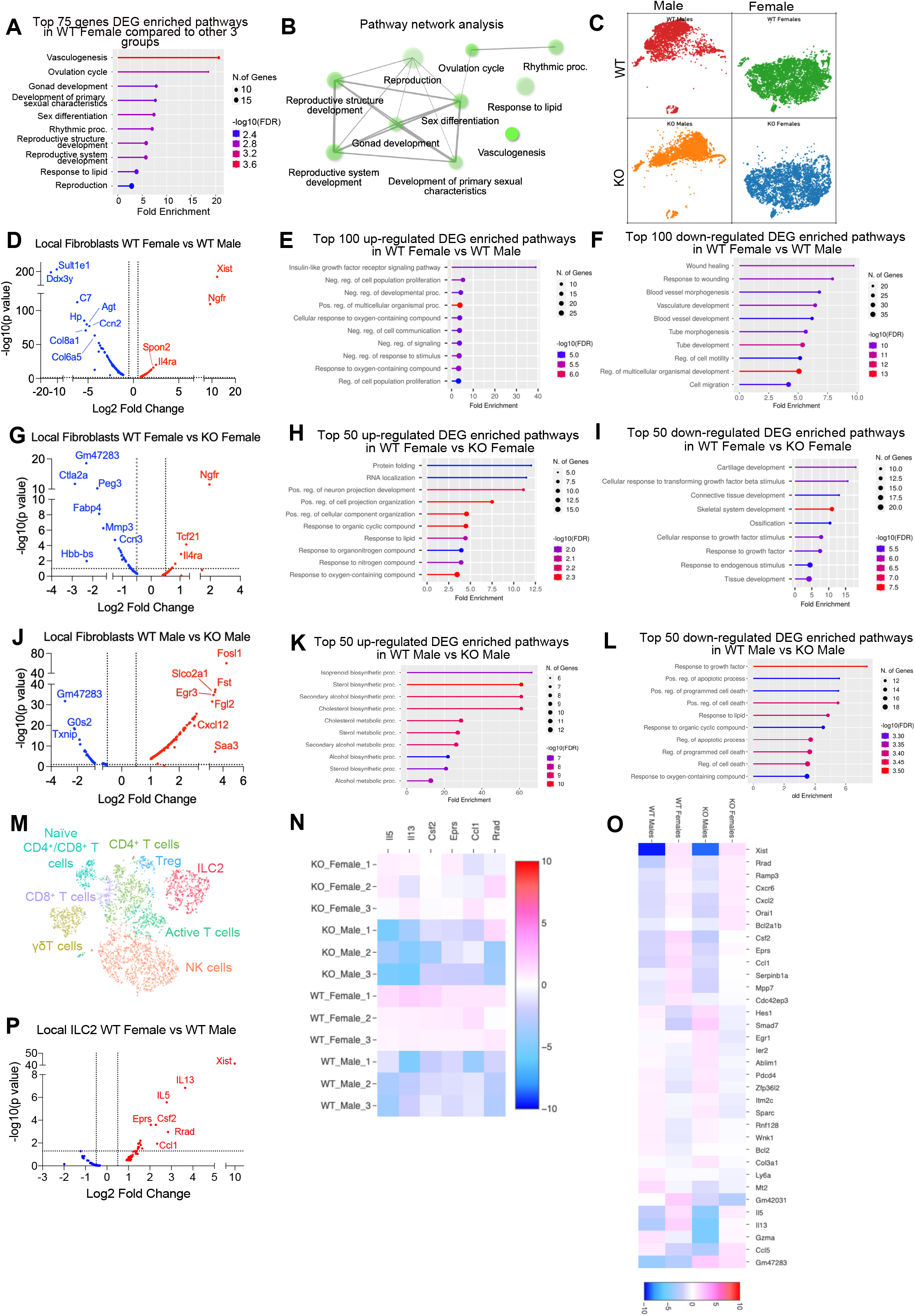
Differentially expressed genes in fibroblast and ILC2 cell populations from scRNA-seq analysis. **(A)** Gene ontology (GO) pathways of the top 75 genes that were upregulated in WT females vs other three group in all clusters. **(B)** Pathway network analysis of the top 75 genes upregulated in WT females vs other three groups. **(C)** Population-shift in fibroblasts between WT female, WT male, KO female and KO male. **(D-L)** Differentially expressed genes in fibroblast population between WT female vs. WT male, WT female vs KO female, WT male vs KO male as associated GO terms for top DEG. **(M-P)** ILC2 cell clusters and DEG between WT females and WT males. Heatmaps indicate DEGs between WT female, WT male, KO female and KO males.

**Figure S3.**
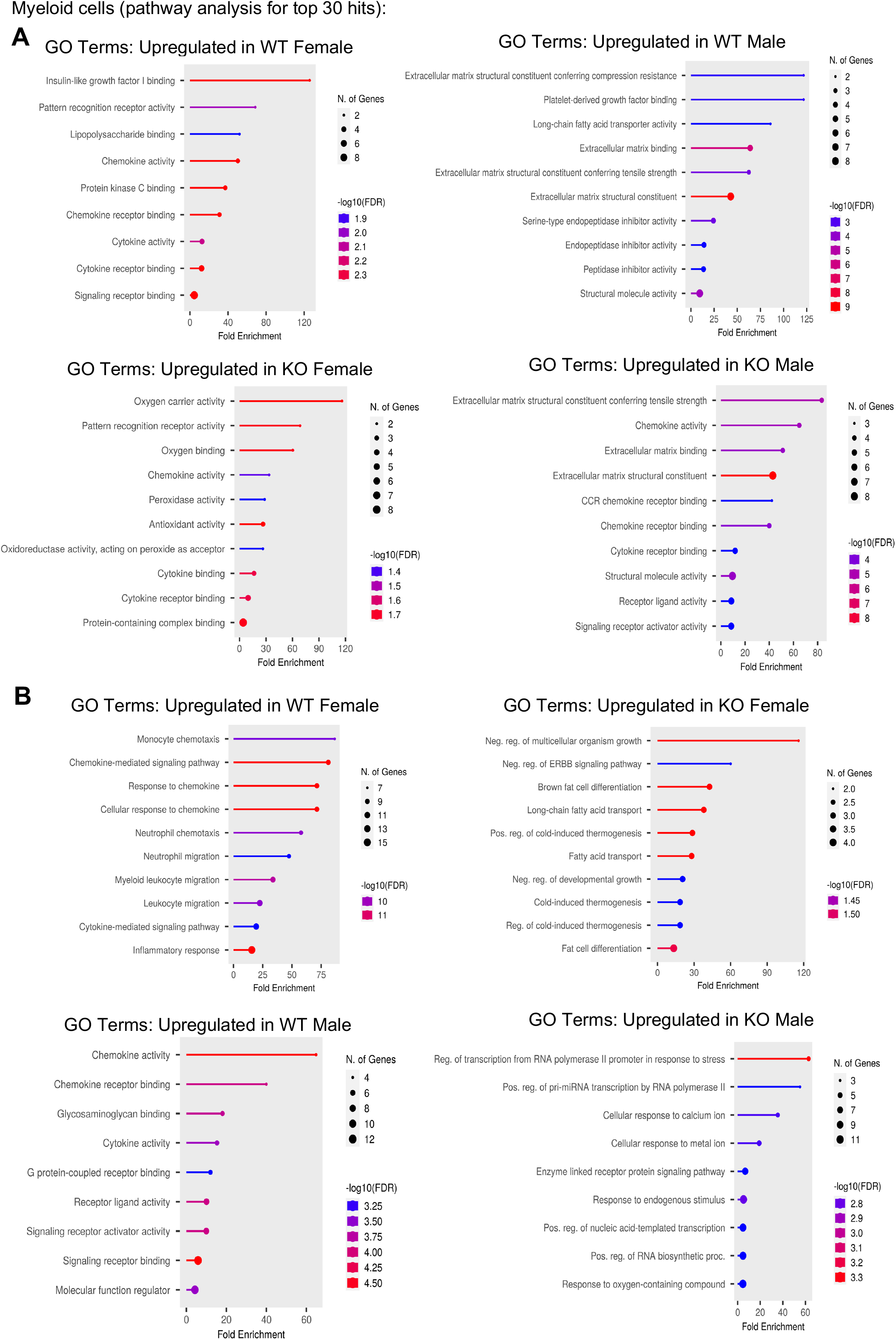
Gene ontology pathway analysis for the top 30 hits in myeloid cell subsets. **(A)** GO pathway analyses comparing the top 30 genes between WT female vs WT male, and KO female vs KO male in myeloid cell subsets. **(B)**: GO comparing the top 30 genes between WT female vs KO female and WT male vs KO male in myeloid cell subsets

**Figure S4.**
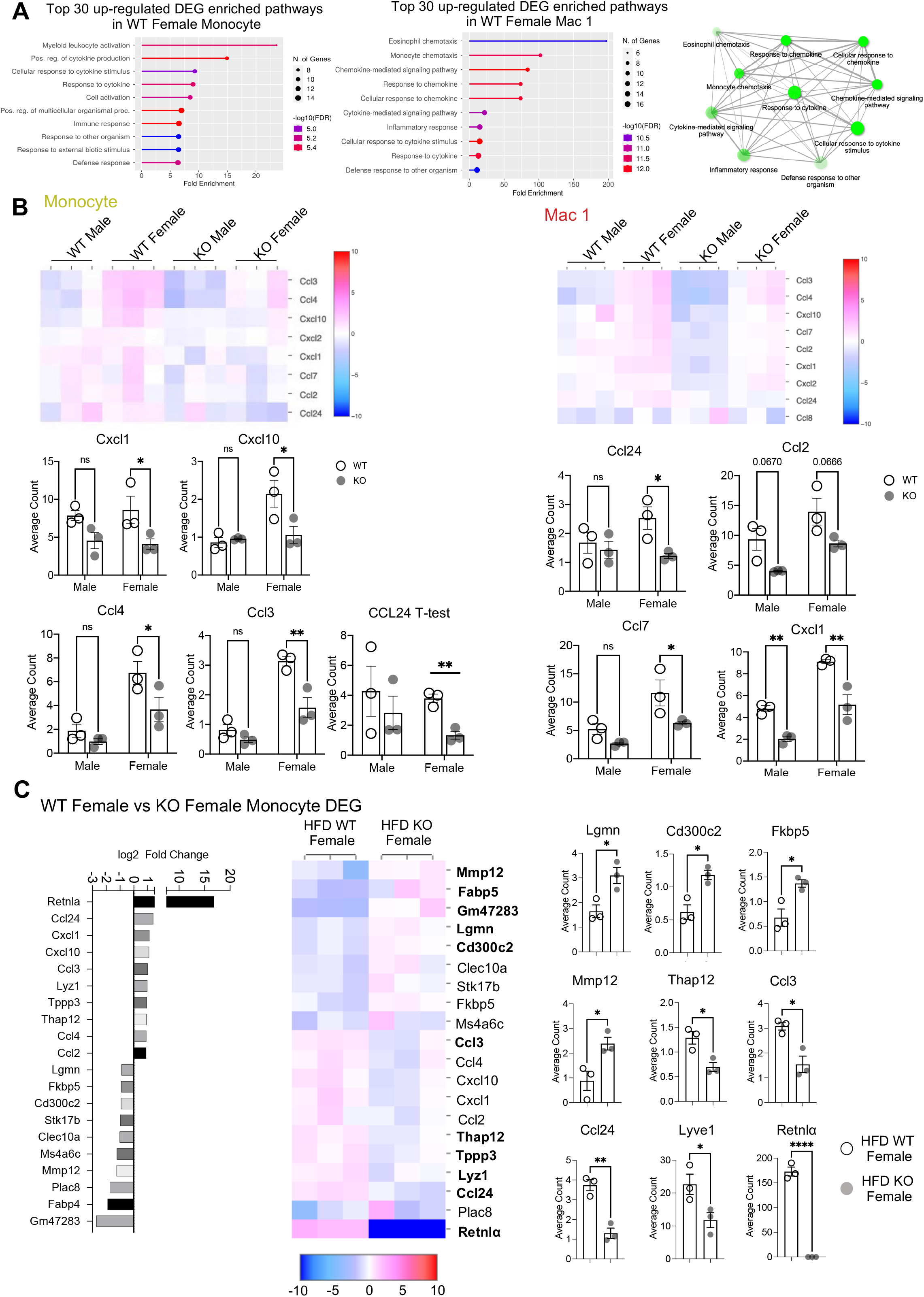
Monocyte and Mac1 DEG. **(A)** GO pathway analyses of top 30 DEG in WT female monocytes and Mac1 compared to other groups. **(B)** Heatmaps indicating DEGs in Monocyte and Mac1 and average UMI count analysis between all groups for key marker genes. **(C)** WT female vs KO female on HFD monocyte subcluster DEG presented as Log2 fold changes in a heatmap and average UMI count comparisons.

**Supplemental Table 1.** Three-way ANOVA analysis of Body weight, Immune cells, Macrophages and Eosinophils.

Mouse body weight 6 weeks Post Diet **3Way ANOVA**
| Source of Variation | % of total variation | P value | P value summary | Significant? |
| --- | --- | --- | --- | --- |
| Male Vs Female | 34.40 | <0.0001 | **** | Yes |
| Ctr Vs HFD | 34.04 | <0.0001 | **** | Yes |
| WT Vs KO | 4.226 | 0.0056 | ** | Yes |
| Male Vs Female x Ctr Vs HFD | 5.005 | 0.0028 | ** | Yes |
| Male Vs Female x WT Vs KO | 2.019 | 0.0484 | * | Yes |
| Ctr Vs HFD x WT Vs KO | 0.6178 | 0.2651 | ns | No |
| Male Vs Female x Ctr Vs HFD x WT Vs KO | 3.861 | 0.0078 | ** | Yes |

Mouse body weight 12 weeks Post Diet **3Way ANOVA**
| Source of Variation | % of total variation | P value | P value summary | Significant? |
| --- | --- | --- | --- | --- |
| Male Vs Female | 13.13 | <0.0001 | **** | Yes |
| Ctr Vs HFD | 71.83 | <0.0001 | **** | Yes |
| WT Vs KO | 3.335 | 0.0003 | *** | Yes |
| Male Vs Female x Ctr Vs HFD | 1.000 | 0.0319 | * | Yes |
| Male Vs Female x WT Vs KO | 1.668 | 0.0067 | ** | Yes |
| Ctr Vs HFD x WT Vs KO | 2.663 | 0.0009 | *** | Yes |
| Male Vs Female x Ctr Vs HFD x WT Vs KO | 1.668 | 0.0067 | ** | Yes |

Leukocytes in % of SVF **3Way ANOVA**
| Source of Variation | % of total variation | P value | P value summary | Significant? |
| --- | --- | --- | --- | --- |
| Male Vs Female | 9.708 | <0.0001 | **** | Yes |
| Ctr Vs HFD | 55.05 | <0.0001 | **** | Yes |
| WT Vs KO | 17.48 | <0.0001 | **** | Yes |
| Male Vs Female x Ctr Vs HFD | 3.345 | 0.0043 | ** | Yes |
| Male Vs Female x WT Vs KO | 0.005690 | 0.8980 | ns | No |
| Ctr Vs HFD x WT Vs KO | 0.3134 | 0.3459 | ns | No |
| Male Vs Female x Ctr Vs HFD x WT Vs KO | 7.520 | <0.0001 | **** | Yes |

Macrophage in % of Leukocytes **3Way ANOVA**
| Source of Variation | % of total variation | P value | P value summary | Significant? |
| --- | --- | --- | --- | --- |
| Male VS Female | 8.384 | 0.0068 | ** | Yes |
| Ctr Vs HFD | 17.40 | 0.0003 | *** | Yes |
| WT Vs KO | 2.665 | 0.1084 | ns | No |
| Male VS Female x Ctr Vs HFD | 38.38 | <0.0001 | **** | Yes |
| Male VS Female x WT Vs KO | 0.6558 | 0.4167 | ns | No |
| Ctr Vs HFD x WT Vs KO | 7.337 | 0.0106 | * | Yes |
| Male VS Female x Ctr Vs HFD x WT Vs KO | 0.0003325 | 0.9853 | ns | No |

Eosinophils in % of Leukocytes **3Way ANOVA**
| Source of Variation | % of total variation | P value | P value summary | Significant? |
| --- | --- | --- | --- | --- |
| Male Vs Female | 54.11 | <0.0001 | **** | Yes |
| Ctr Vs HFD | 19.61 | <0.0001 | **** | Yes |
| WT Vs KO | 1.364 | 0.1743 | ns | No |
| Male Vs Female x Ctr Vs HFD | 2.298 | 0.0815 | ns | No |
| Male Vs Female x WT Vs KO | 0.001983 | 0.9579 | ns | No |
| Ctr Vs HFD x WT Vs KO | 1.396 | 0.1695 | ns | No |
| Male Vs Female x Ctr Vs HFD x WT Vs KO | 3.595 | 0.0321 | * | Yes |

## References

1. H. The Lancet Public, Tackling obesity seriously: the time has come. The Lancet. Public health 3, e153 (2018).

2. J. C. Link, K. Reue, Genetic Basis for Sex Differences in Obesity and Lipid Metabolism. Annual review of nutrition 37, 225–245 (2017).

3. E. Gerdts, V. Regitz-Zagrosek, Sex differences in cardiometabolic disorders. Nature medicine 25, 1657–1666 (2019).

4. B. W. Parks et al., Genetic architecture of insulin resistance in the mouse. Cell metabolism 21, 334–347 (2015).

5. J. P. Camporez et al., Anti-inflammatory effects of oestrogen mediate the sexual dimorphic response to lipid-induced insulin resistance. The Journal of physiology 597, 3885–3903 (2019).

6. K. E. Chen, N. M. Lainez, D. Coss, Sex Differences in Macrophage Responses to Obesity-Mediated Changes Determine Migratory and Inflammatory Traits. Journal of immunology 206, 141–153 (2021).

7. N. M. Lainez et al., Diet-Induced Obesity Elicits Macrophage Infiltration and Reduction in Spine Density in the Hypothalami of Male but Not Female Mice. Frontiers in immunology 9, 1992 (2018).

8. B. F. Palmer, D. J. Clegg, The sexual dimorphism of obesity. Mol Cell Endocrinol 402, 113–119 (2015).

9. B. L. Wajchenberg, Subcutaneous and visceral adipose tissue: their relation to the metabolic syndrome. Endocrine reviews 21, 697–738 (2000).

10. S. P. Weisberg et al., Obesity is associated with macrophage accumulation in adipose tissue. The Journal of clinical investigation 112, 1796–1808 (2003).

11. C. A. Curat et al., Macrophages in human visceral adipose tissue: increased accumulation in obesity and a source of resistin and visfatin. Diabetologia 49, 744–747 (2006).

12. J. M. Olefsky, C. K. Glass, Macrophages, inflammation, and insulin resistance. Annu Rev Physiol 72, 219–246 (2010).

13. M. L. Gruen, M. Hao, D. W. Piston, A. H. Hasty, Leptin requires canonical migratory signaling pathways for induction of monocyte and macrophage chemotaxis. American Journal of Physiology-Cell Physiology 293, C1481–C1488 (2007).

14. H. Kanda et al., MCP-1 contributes to macrophage infiltration into adipose tissue, insulin resistance, and hepatic steatosis in obesity. The Journal of clinical investigation 116, 1494–1505 (2006).

15. J. L. Kaplan et al., Adipocyte progenitor cells initiate monocyte chemoattractant protein-1-mediated macrophage accumulation in visceral adipose tissue. Molecular metabolism 4, 779–794 (2015).

16. D. E. Lackey, J. M. Olefsky, Regulation of metabolism by the innate immune system. Nat Rev Endocrinol 12, 15–28 (2016).

17. J. C. McNelis, J. M. Olefsky, Macrophages, immunity, and metabolic disease. Immunity 41, 36–48 (2014).

18. S. T. Gal-Oz et al., ImmGen report: sexual dimorphism in the immune system transcriptome. Nat Commun 10, 4295 (2019).

19. M. Varghese et al., Monocyte Trafficking and Polarization Contribute to Sex Differences in Meta-Inflammation. Frontiers in endocrinology 13, 826320 (2022).

20. K. J. Strissel et al., Adipocyte death, adipose tissue remodeling, and obesity complications. Diabetes 56, 2910–2918 (2007).

21. K. E. Chen, N. M. Lainez, M. G. Nair, D. Coss, Visceral adipose tissue imparts peripheral macrophage influx into the hypothalamus. Journal of neuroinflammation 18, 140 (2021).

22. A. Keselman, X. Fang, P. B. White, N. M. Heller, Estrogen Signaling Contributes to Sex Differences in Macrophage Polarization during Asthma. Journal of immunology 199, 1573–1583 (2017).

23. K. L. Grove, S. K. Fried, A. S. Greenberg, X. Q. Xiao, D. J. Clegg, A microarray analysis of sexual dimorphism of adipose tissues in high-fat-diet-induced obese mice. International journal of obesity (2005) 34, 989–1000 (2010).

24. R. E. Stubbins, V. B. Holcomb, J. Hong, N. P. Nunez, Estrogen modulates abdominal adiposity and protects female mice from obesity and impaired glucose tolerance. European journal of nutrition 51, 861–870 (2012).

25. E. L. Sullivan, A. J. Daniels, F. H. Koegler, J. L. Cameron, Evidence in female rhesus monkeys (Macaca mulatta) that nighttime caloric intake is not associated with weight gain. Obesity research 13, 2072–2080 (2005).

26. P. A. Heine, J. A. Taylor, G. A. Iwamoto, D. B. Lubahn, P. S. Cooke, Increased adipose tissue in male and female estrogen receptor-alpha knockout mice. Proceedings of the National Academy of Sciences of the United States of America 97, 12729–12734 (2000).

27. B. B. Shenoda et al., Xist attenuates acute inflammatory response by female cells. Cellular and molecular life sciences: CMLS, (2020).

28. K. Singer et al., Differences in Hematopoietic Stem Cells Contribute to Sexually Dimorphic Inflammatory Responses to High Fat Diet-induced Obesity. The Journal of biological chemistry 290, 13250–13262 (2015).

29. S. Chakarov, C. Blériot, F. Ginhoux, Role of adipose tissue macrophages in obesity-related disorders. The Journal of experimental medicine 219, (2022).

30. D. Wu et al., Eosinophils sustain adipose alternatively activated macrophages associated with glucose homeostasis. Science 332, 243–247 (2011).

31. J. R. Brestoff et al., Group 2 innate lymphoid cells promote beiging of white adipose tissue and limit obesity. Nature 519, 242–246 (2015).

32. D. A. Hill et al., Distinct macrophage populations direct inflammatory versus physiological changes in adipose tissue. Proceedings of the National Academy of Sciences 115, e5096–E5105 (2018).

33. D. A. Jaitin et al., Lipid-Associated Macrophages Control Metabolic Homeostasis in a Trem2-Dependent Manner. Cell 178, 686–698.e614 (2019).

34. G. M. Pine, H. M. Batugedara, M. G. Nair, Here, there and everywhere: Resistin-like molecules in infection, inflammation, and metabolic disorders. Cytokine 110, 442–451 (2018).

35. M. Lv, W. Liu, Hypoxia-Induced Mitogenic Factor: A Multifunctional Protein Involved in Health and Disease. Frontiers in cell and developmental biology 9, 691774 (2021).

36. J. Li, S. Y. Kim, N. M. Lainez, D. Coss, M. G. Nair, Macrophage-Regulatory T Cell Interactions Promote Type 2 Immune Homeostasis Through Resistin-Like Molecule α. Frontiers in immunology 12, 710406 (2021).

37. T. A. Harris et al., Resistin-like Molecule α Provides Vitamin-A-Dependent Antimicrobial Protection in the Skin. Cell host & microbe 25, 777–788.e778 (2019).

38. B. Krljanac et al., RELMα-expressing macrophages protect against fatal lung damage and reduce parasite burden during helminth infection. Science immunology 4, (2019).

39. D. E. Sanin et al., A common framework of monocyte-derived macrophage activation. Science immunology 7, eabl7482 (2022).

40. C. C. Bain et al., CD11c identifies microbiota and EGR2-dependent MHCII(+) serous cavity macrophages with sexually dimorphic fate in mice. European journal of immunology 52, 1243–1257 (2022).

41. Y. Kumamoto et al., CD301b(+) Mononuclear Phagocytes Maintain Positive Energy Balance through Secretion of Resistin-like Molecule Alpha. Immunity 45, 583–596 (2016).

42. M. R. Lee et al., The adipokine Retnla modulates cholesterol homeostasis in hyperlipidemic mice. Nat Commun 5, 4410 (2014).

43. A. E. Salinero, B. M. Anderson, K. L. Zuloaga, Sex differences in the metabolic effects of diet-induced obesity vary by age of onset. International journal of obesity (2005) 42, 1088–1091 (2018).

44. F. Chen et al., B Cells Produce the Tissue-Protective Protein RELMα during Helminth Infection, which Inhibits IL-17 Expression and Limits Emphysema. Cell Rep 25, 2775–2783.e2773 (2018).

45. L. Liu, M. Zeng, J. S. Stamler, Hemoglobin induction in mouse macrophages. Proceedings of the National Academy of Sciences of the United States of America 96, 6643–6647 (1999).

46. D. Saha et al., Hemoglobin Expression in Nonerythroid Cells: Novel or Ubiquitous? International Journal of Inflammation 2014, 803237 (2014).

47. A. Weinstock, H. Moura Silva, K. J. Moore, A. M. Schmidt, E. A. Fisher, Leukocyte Heterogeneity in Adipose Tissue, Including in Obesity. Circulation research 126, 1590–1612 (2020).

48. M. Ikutani, S. Nakae, Heterogeneity of Group 2 Innate Lymphoid Cells Defines Their Pleiotropic Roles in Cancer, Obesity, and Cardiovascular Diseases. Frontiers in immunology 13, 939378 (2022).

49. M. W. Lee et al., Activated type 2 innate lymphoid cells regulate beige fat biogenesis. Cell 160, 74–87 (2015).

50. A. Munitz et al., Resistin-like molecule alpha decreases glucose tolerance during intestinal inflammation. Journal of immunology 182, 2357–2363 (2009).

51. G. Vlacich, M. C. Nawijn, G. C. Webb, D. F. Steiner, Pim3 negatively regulates glucose-stimulated insulin secretion. Islets 2, 308–317 (2010).

52. M. P. Godard, B. A. Johnson, S. R. Richmond, Body composition and hormonal adaptations associated with forskolin consumption in overweight and obese men. Obesity research 13, 1335–1343 (2005).

53. N. Mukaida, Y. Y. Wang, Y. Y. Li, Roles of Pim-3, a novel survival kinase, in tumorigenesis. Cancer science 102, 1437–1442 (2011).

54. W. G. Tharp et al., Effects of glucose and insulin on secretion of amyloid-β by human adipose tissue cells. Obesity (Silver Spring, Md.) 24, 1471–1479 (2016).

55. Z. Wan, D. Mah, S. Simtchouk, A. Kluftinger, J. P. Little, Role of amyloid β in the induction of lipolysis and secretion of adipokines from human adipose tissue. Adipocyte 4, 212–216 (2015).

56. C. A. Ihunnah et al., Estrogen sulfotransferase/SULT1E1 promotes human adipogenesis. Molecular and cellular biology 34, 1682–1694 (2014).

57. J. M. Castellano, J. M. Espinosa, J. S. Perona, Modulation of Lipid Transport and Adipose Tissue Deposition by Small Lipophilic Compounds. Frontiers in cell and developmental biology 8, (2020).

58. F. Gao et al., Suppression of lncRNA Gm47283 attenuates myocardial infarction via miR-706/ Ptgs2/ferroptosis axis. Bioengineered 13, 10786–10802 (2022).

59. Y. Qiu et al., Eosinophils and type 2 cytokine signaling in macrophages orchestrate development of functional beige fat. Cell 157, 1292–1308 (2014).

60. W. R. Bolus et al., Elevating adipose eosinophils in obese mice to physiologically normal levels does not rescue metabolic impairments. Molecular metabolism 8, 86–95 (2018).

61. C. L. Grek, D. A. Newton, D. D. Spyropoulos, J. E. Baatz, Hypoxia up-regulates expression of hemoglobin in alveolar epithelial cells. American journal of respiratory cell and molecular biology 44, 439–447 (2011).

62. D. A. Gell, Structure and function of haemoglobins. Blood cells, molecules & diseases 70, 13–42 (2018).

63. M. Orecchioni, Y. Ghosheh, A. B. Pramod, K. Ley, Macrophage Polarization: Different Gene Signatures in M1(LPS+) vs. Classically and M2(LPS-) vs. Alternatively Activated Macrophages. Frontiers in immunology 10, 1084 (2019).

64. A. C. Cepeda-Lopez, K. Baye, Obesity, iron deficiency and anaemia: a complex relationship. Public Health Nutrition 23, 1703–1704 (2020).

65. C. C. Bain et al., Rate of replenishment and microenvironment contribute to the sexually dimorphic phenotype and function of peritoneal macrophages. Science immunology 5, (2020).

66. D. E. Sanin et al., A common framework of monocyte-derived macrophage activation. Science immunology 7, eabl7482 (2022).

67. S. X. Ge, D. Jung, R. Yao, ShinyGO: a graphical gene-set enrichment tool for animals and plants. Bioinformatics (Oxford, England) 36, 2628–2629 (2020).

68. X. Qiu et al., Reversed graph embedding resolves complex single-cell trajectories. Nature methods 14, 979–982 (2017).

69. L. Haghverdi, A. T. L. Lun, M. D. Morgan, J. C. Marioni, Batch effects in single-cell RNA-sequencing data are corrected by matching mutual nearest neighbors. Nature biotechnology 36, 421–427 (2018).

